# Protein prediction models support widespread post-transcriptional regulation of protein abundance by interacting partners

**DOI:** 10.1101/2022.03.14.484316

**Authors:** Himangi Srivastava, Michael J. Lippincott, Jordan Currie, Robert Canfield, Maggie P. Y. Lam, Edward Lau

## Abstract

Protein and mRNA levels correlate only moderately. The availability of proteogenomics data sets with protein and transcript measurements from matching samples is providing new opportunities to assess the degree to which protein levels in a system can be predicted from mRNA information. Here we examined the contributions of input features in protein abundance prediction models. Using large proteogenomics data from 8 cancer types within the Clinical Proteomic Tumor Analysis Consortium (CPTAC) data set, we trained models to predict the abundance of over 13,000 proteins using matching transcriptome data from up to 958 tumor or normal adjacent tissue samples each, and compared predictive performances across algorithms, data set sizes, and input features. Over one-third of proteins (4,648) showed relatively poor predictability (elastic net r ≤ 0.3) from their cognate transcripts. Moreover, we found widespread occurrences where the abundance of a protein is considerably less well explained by its own cognate transcript level than that of one or more trans locus transcripts. The incorporation of additional trans-locus transcript abundance data as input features increasingly improved the ability to predict sample protein abundance. Transcripts that contribute to non-cognate protein abundance primarily involve those encoding known or predicted interaction partners of the protein of interest, including not only large multi-protein complexes as previously shown, but also small stable complexes in the proteome with only one or few stable interacting partners. Network analysis further shows a complex proteome-wide interdependency of protein abundance on the transcript levels of multiple interacting partners. The predictive model analysis here therefore supports that protein-protein interaction including in small protein complexes exert post-transcriptional influence on proteome compositions more broadly than previously recognized. Moreover, the results suggest mRNA and protein co-expression analysis may have utility for finding gene interactions and predicting expression changes in biological systems.

## Introduction

Mounting evidence now shows that protein levels correlate imperfectly with the levels of their cognate transcripts ^1–3^. More specifically, although a robust trend exists over the log scale between protein and mRNA measurements *across genes*, genewise correlation between proteins and their transcripts is much poorer *across observations* (samples, tissues, cell types, or subjects). This has been taken to indicate that while abundant proteins have abundant transcripts, transcript variance within a group of samples does not necessarily predict or signify corresponding protein changes ^4^. Multiple factors are known to contribute to this non-correlation. Technical variations are often cited as a substantial source of non-correlation, as transcriptomics and proteomics measurements carry different sources of error and proteins with lower baseline variance in mass spectrometry have been shown to be better predicted by their transcripts ^5^. Nevertheless, a substantial portion of protein variance remains unexplained and is likely attributable to biological and biophysical regulations. It has been well recognized that large multi-protein complexes could invoke a buffer effect on protein levels ^6,7^, as a multimeric complex only fully folds and functions when all subunits are present, any induction of the transcript for a single subunit would not per se lead to additional complexes, and resulting supernumerary proteins are thought to be quickly degraded ^6,8^. Lastly, numerous post-transcriptional and post-translational mechanisms are known to modulate protein levels such as the gene- and context-dependent translation rates of mRNAs ^9,10^, the differential half-life and temporal distributions between mRNAs and proteins ^2, 11^, and proteolytic degrading translated proteins in the cell.

The emergence of large-scale proteogenomics data from matching samples have created new opportunities to revisit protein-level predictions from transcriptomics data. The abundance of a protein may be the function of one or more transcripts. Most notably, available data sets from the Gene Tissue Expression (GTEx) project ^7^ and the Clinical Proteomic Tumor Analysis Consortium (CPTAC) ^12^, have spurred the use of machine learning approaches to evaluate how well one can predict protein level variance given a set of transcriptomics data, with the goal of developing strategies and algorithms that can boost the performance of protein level predictions. This culminated in a community based effort in the CPTAC Proteogenomics Dream Challenge Task 2, which tasked participants with predicting protein abundances from mRNA and genetic data from CPTAC ovarian and breast cancer samples ^12^. The results suggest that protein level prediction remains a challenging and not fully resolved problem, as many community submitted models did not improve substantially the baseline model, which is an elastic net taking into account all mRNA features available and has a median Pearson’s correlation coefficient (*r*) of 0.47 for ovarian cancer. Nevertheless, general lessons have emerged from the top performing models; for instance: (i) ensemble methods generally performed well ^12–14^; (ii) combining observations from the ovarian and breast cancer datasets to borrow information from each other led to improved predictions ^12^; and (iii) judicious feature preselection based on prior biological knowledge such as protein-protein interactions improved prediction performance ^12,14^. Notwithstanding these general observations, the current literature reflects that much remains to be learned about the relationship of mRNA and protein regulations in different genes and whether there are fundamental limits to how well mRNA abundance reflects that of their protein counterpart. This problem has several practical importances. Proteins carry out the majority of biological processes and hence are arguably the most relevant molecules to biological states. Despite rapid advances in proteomics techniques, bulk and single-cell RNA sequencing remain the most commonly used methods to interrogate gene expression status on a large scale and will likely remain so in the foreseeable future. Transcriptomics experiments often operate on the implicit assumption that identified differential regulation exert their biological effects via their cognate proteins, hence it is important to better understand the relationships between protein and mRNA levels to aid in data interpretation and determining potential protein level changes given a set of transcriptomics data. Alternatively, knowing the genewise difference in how well a gene’s transcript can predict its protein counterpart may be useful for filtering and prioritizing biologically relevant transcript signatures ^15^.

Here we revisit the predictability of protein levels from transcriptomics data. Since the time of the Dream Challenge, considerably more proteogenomics data have been made publicly available which increases the number of observations available for modeling training, as well as the number of proteins for which there is mass spectrometry information available. Individual CPTAC cancer studies have analyzed the protein and mRNA correlation in individual tumors and normal adjacent tissues and nominated specific pathways whose correlations are particularly poor. Additional re-analysis and meta-analysis studies have outlined the distribution of prediction performances across algorithms, and generally conclude there is some statistical enrichment of biological processes or protein features among proteins that are poorly predicted by their own transcript level, e.g., metabolic and essential proteins or proteins belonging to complexes ^12,16^. Nevertheless, a granular analysis remains unrealized in the literature that interrogates the identity and regulatory modality of individual proteins in depth. Accordingly, our goals here are to (1) evaluate how the increasing data size from combining CPTAC tumor data sets affects the performance of prediction algorithms and feature selection strategies; and (2) interpret prediction models to assess the importance of transcript features in individual protein abundance regulation. The results suggest that the incorporation of transcript level information from protein interacting partners played a substantial role in predicting protein levels, and moreover, there are widespread instances in the proteome where the abundance of a protein correlates primarily with a trans locus transcript than its own cognate transcript, which has implications for gene expression profiling studies.

## Methods

### Data retrieval and processing

Gene expression data were obtained from public data from the CPTAC project and included data from 8 cancer types: ovarian cancer (OV) ^17^, breast cancer (BR) ^18^, endometrial carcinoma (EN) ^19^, colorectal cancer (CO) ^20^, lung adenocarcinoma (LUAD) ^21^, clear cell renal carcinoma (CCRCC) ^22^, glioblastoma (GB) ^23^, and lung squamous cell carcinoma (LSCC) ^24^. The cumulative inclusions of each cancer type in the order above are sequentially referred to as CPTAC_2 to CPTAC_8 in the manuscript, such that CPTAC_2 refers to the union of ovarian and breast cancer (OV + BR); CPTAC_3 refers the union of ovarian, breast, and endometrial cancer (OV + BR + EN); and so on. The mRNA and protein level expression data from the CPTAC cancer types was retrieved using the cptac package v.0.9.7 ^25^ in Python 3.9. Each column of the quantitative measurement of the transcriptomics data acted as an independent variable or feature variable whereas the normalized quantitative measurement of a particular protein of interest acted as the single dependent or target variable in the protein model. Retrieved mRNA level gene expression data are standardized using the scikit-learn simple scaler. The proteomics data were likewise downloaded using the cptac package as presented in the data, and were stable isotope labeled relative quantitative mass spectrometry data presented as normalized log ratios across samples as in the original studies. All tumor samples were labeled using Thermo tandem mass tag (TMT) 10- or 11- plex isobaric tags for MS2 quantification, with the exception of the ovarian cancer data, which were labeled with Sciex iTRAQ isobaric tags for MS2 quantification, and the colon cancer data, which contained both label-free and TMT quantifications. The retrieved log ratios across samples were not further transformed.

For each protein for which predictions are to be made, we retrieved five separate feature sets:

1. **Single:** Using only the single transcript coding for the protein of interest for model training, then running the pipeline to train a model for each protein separately.
2. **CORUM:** Using the transcripts of all proteins belonging to the same protein complex as the protein of interest, if any, in CORUM v.3.0 ^26^, where protein complexes are defined as two or more proteins that interact physically in a quaternary structure. These transcripts then act as independent variables (features) to predict the target variable (protein of interest). The pipeline is then run to train a model for each protein separately.
3. **STRING 800:** Using the transcripts of interacting partners of the protein of interest as input features. Interacting partners are retrieved from STRING v.11 ^27^, which documents functional associations including physical interactions, genetic interactions, co-expression, co-occurrence, and other associations. The STRING combined score represents the overall likelihood of interactions. Interacting pairs with a STRING combined score of 800 or above (high-confidence) are included. The pipeline is then run to train a model for each protein separately.
4. **STRING 200:** As above, except that interacting pairs with a STRING combined score of 200 or above (low- to high-confidence) are included. The pipeline is then run to train a model for each protein separately.
5. **Transcriptome:** A transcriptome-wide model where all qualifying transcripts in the data set are included as features, prior to the removal of low-variance features. The pipeline is then run to train a model for each protein separately.

### Model training and evaluation

For each feature set, missing values for feature variables are imputed using median imputation, followed by the removal of features with variance of 0.2 or below. Models are not trained for proteins with fewer than 50 empirical observations. The data are then split 80:20 into training and test sets. No imputation or additional standardization was performed on the proteomics data. The input and target data from the training set are then used to train a model using either linear regression, elastic net with 5-fold cross validation, random forest regressor, or gradient boosting regressor in scikit-learn v.1.0 ^28^, with the following specified parameters: random forest number of estimators: 500, criterion: squared error, max depth: 4; elastic net cross validation L1 ratio: 0.1, 0.5, 0.9, 0.95, cv: 5, tolerance: 1e–3, max iterations: 2000; gradient boosting: n_estimators: 1000, max_depth: 3, subsample: 0.5, min_samples_splot: 5, learning_rate: 0.025. The trained models are saved as individual objects which include the predicted protein levels in the training set as well as the contribution of each feature to the overall prediction (coefficients in linear regression and elastic net; feature importance and trees in random forest and gradient boosting regressors), and are applied to predict protein levels in the test set data. Further interpretation of feature importance was performed using Shapley values with the aid of the shap package v.0.40.0 ^29^, or with the Boruta algorithm using the Boruta_Py package v.0.3 ^30^.

To evaluate model performance, the Pearson’s correlation coefficients between predicted 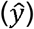 and actual (*y*) protein values are calculated individually for the train and test set data for each protein model using the numpy corrcoef function, or defaulted to 0 if the standard deviation of predicted values is 0. Goodness-of-fit (R^2^) is calculated using scikit-learn. metrics. r2_score function. Normalized root mean square errors (NRMSE) are calculated as follows:

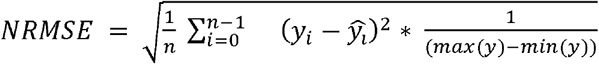

### Network construction and analysis

To construct the protein regulation network, we first constructed an overall graph from a subset of transcript-protein relationships using the networkx package ^31^ in Python 3.9. Qualifying proteins are those whose prediction increased with the inclusion of more features, such that in the STRING or CORUM feature set, the correlation coefficient between the elastic net predicted and actual protein levels is greater than the single feature elastic nets by at least 0.25. Transcripts that contribute to the prediction of these proteins are therefore included if their elastic net coefficient *a* and random forest feature importance *b* are greater than a certain threshold, which was set to *a*=0.05, *b*=0.05 for the CORUM feature set and *a*=0.2, *b*=0.05 for the STRING feature set.

In the data, the list of proteins and their corresponding transcriptomics features act as the nodes and the interaction pattern is a function of the correlation value and feature importance between a protein and the transcriptomics. The direction of the edge therefore flows from transcriptomics to protein. All the text files for each protein obtained from the computational pipeline are compiled into a n overall data frame. This data frame is then converted into the form of a network data frame. In this data frame, all the protein becomes the target node, and the transcriptomics corresponding to the protein becomes the source node. Additional column of this data frame consists of the weight between a protein and its predicting transcripts, which was calculated using the sum of the [−1,1] clipped elastic net coefficient and the descending percentile rank of the random forest feature importance. An overall directed graph G(V, E) is then constructed from the edge lists with proteins as targets and their predicting transcripts as sources, such that it is a function of vertices V which depict protein and/or transcript nodes and edges E which flow in the direction from the transcript sources to the protein targets. Using the built-in functions in networkx, this overall directed graph is divided into weakly connected components, which resulted in individual subgraphs such that for the overall network G=(V, E) where V is the vertices and E is the edge then a subgraph of G = (V, E) is a graph S=(V’, E’) where vertex set V’ ⊆ V and edge set E’ ⊆ E connects only nodes of V’. We then used Cytoscape v.3.9 ^32^ to visualize and perform additional topology analysis and functional annotations of each subgraph. HiDef persistent community detection ^33^ was performed with the aid of the CyCommunityDetection Cytoscape plugin ^34^ using the Leiden algorithm ^35^. Functional enrichment of individual communities was performed using CyCommunityDetection with the g:Profiler web server ^36^. Hub nodes were identified with the aid of the CytoNCA ^37^ Cytoscape plugin using the betweenness centrality algorithm with the top 10% of proteins with the highest centrality defined as hubs. Functional enrichment analysis on the hub proteins was performed with the aid of the stringApp ^38^ Cytoscape plugin using default settings and 5% false discovery rate cutoff.

### Additional data analysis

Additional data analysis, statistics, and visualization were performed in R v.4.1.1 with the aid of the ReactomePA ^39^, clusterProfiler ^40^, and circlize ^41^ packages; and in Python 3.9 with the aid of the seaborn ^42^ package.

### Data and code availability

Code to train machine learning models and evaluate prediction performance have been uploaded to GitHub at https://github.com/Lau-Lab/CPTAC_Protein. Code for making figures can be found under notebook/PLOS_MakeFig*.qmd.

## Results

### Feature selection improves the prediction of protein abundance from transcriptome data

We first evaluated whether increasing proteogenomics data depth would impact optimal feature selection strategies and algorithms in prediction protein levels. To do so, we retrieved data from up to 8 tumor types from the CPTAC data set and processed the data to collage a data table with matching protein and transcript data from each tumor sample. The data sets were ordered such that breast and ovarian cancers as in the DREAM challenge were combined first, then other cancer types were added according to the order of their availability (see Methods). We trained models to predict the target variable (the normalized labeled mass spectrometry measured abundance of a particular protein) from various input features (transcriptomics data from the matching samples) using either a multiple linear regression, elastic net, and random forest regressor in scikit-learn. To compare the performance of prior knowledge based feature selection, we further compared supplying the models with input features based on (1) only RNA level data the corresponding transcript of the target variable protein (single feature) (2) self transcript plus the transcript of any protein within the same protein complex of the target variable protein in CORUM (CORUM feature); (3) self transcript plus the transcript of any proteins that are high-confidence protein interaction partners including physical interactions and other inferred association from STRINGdb (STRING 800 feature); (4) self transcript plus the transcript of any proteins that are low to high-confidence protein interaction partners from STRINGdb; (5) using all qualifying transcripts (transcriptome features).

We next compared the genewise prediction performance of the transcriptome using the test set correlation coefficient between predicted and actual mass spectrometry protein values, R^2^, and NRMSE as described (**Figure 1A; Supplementary Table S1**). The elastic nets and random fores models performed better with more features and more data. In contrast, a multiple linear regression model performed little better with more included features and in fact failed to predict protein levels when a large number of features are given. For the elastic net and random forest models, performance gains began to saturate with additional data sets (r: 0.388 and 0.402, respectively) but further increased when larger feature sets were given (r: 0.599 for both) (**Figure 1A; Supplementary Figure S1**). As a comparison, we also evaluated the performance of the models in single cancer data (i.e., in each data set alone) (**Supplementary Figure S2A**) as well as combined the data sets in the order of descending performance (**Supplementary Figure S2B**). The single cancer models suggest that a prediction performance of up to median r of 0.691 is achievable using the Transcriptome wide feature set, and 0.69 in the STRING 200 feature set (random forest, CCRCC), albeit in a single cancer type only. In both the single cancer type and reordered combination comparisons, the observation remains true in all cases that the inclusion of protein interactor features further increased the prediction performance of both the elastic net and random forest algorithms. Hence, substantial prediction improvements resulted from the inclusion of other transcripts (number of features) over the gain in data set sizes (number of observations) or the use of algorithms (random forests) that can account for non-linearity. This is corroborated when considering the feature sets across the largest data set collection used (CPTAC_8 from 8 cancer types). Model performance continued to increase as more features were added in (median r from 0.398 to 0.599), and the number of non-predictors and negative predictors decreased while overall dispersion of predictability also decreased.

**Figure 1.**
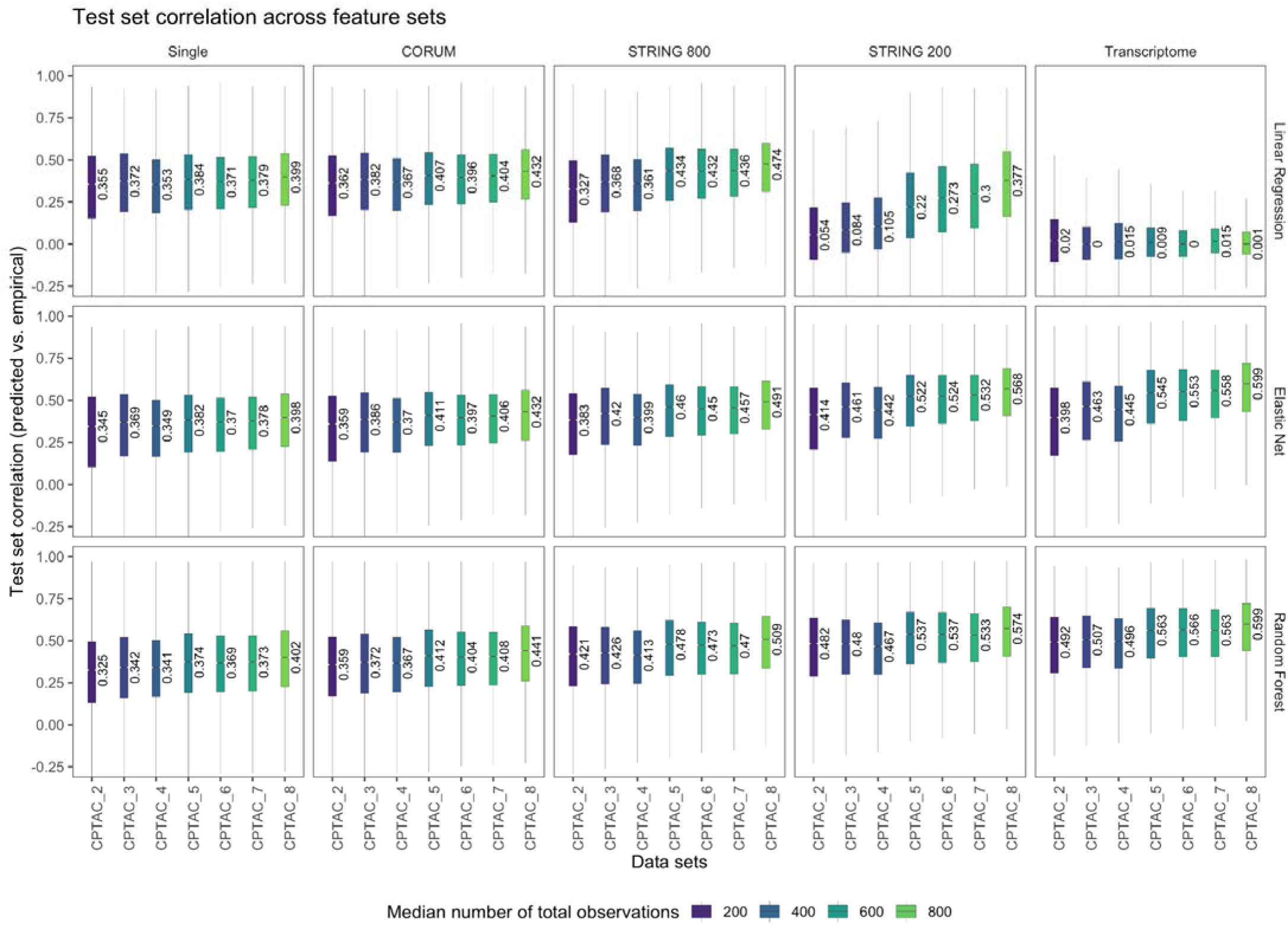
Genewise dispersion of protein predictability from transcriptome data. Box plots of test set correlation coefficients between the transcript-predicted and actual protein level for each protein are shown across five feature sets (column: single/self transcript, CORUM interactors, STRING 800 high-confidence associated proteins; STRING 200 low-confidence associated proteins, and all transcripts) and three algorithms (multiple linear regression, elastic net, and random forest). In each plot, the x axis denotes the number of additive CPTAC data sets used to train the models as described in Methods; box: interquartile range; whiskers: +/− 1.5 IQR; notch: SEM.

From the baseline self-transcript model in the CPTAC_8 data set containing 8 cancer types, we found that strong protein predictors (r ≥ 0.6; 2,008 genes) are enriched in genes participating in membrane proteins and cell junctions (**Figure 2A; Supplementary Table S2**), whereas poor protein predictors (r ≤ 0.3; 4,648 genes) are enriched in multiple large multi-protein complexes (**Figure 2B; Supplementary Table S3**). Hence the analysis of CPTAC_8 data set confirms prior observations that protein complex membership presents a major source for transcript-protein non-correlation.

**Figure 2.**
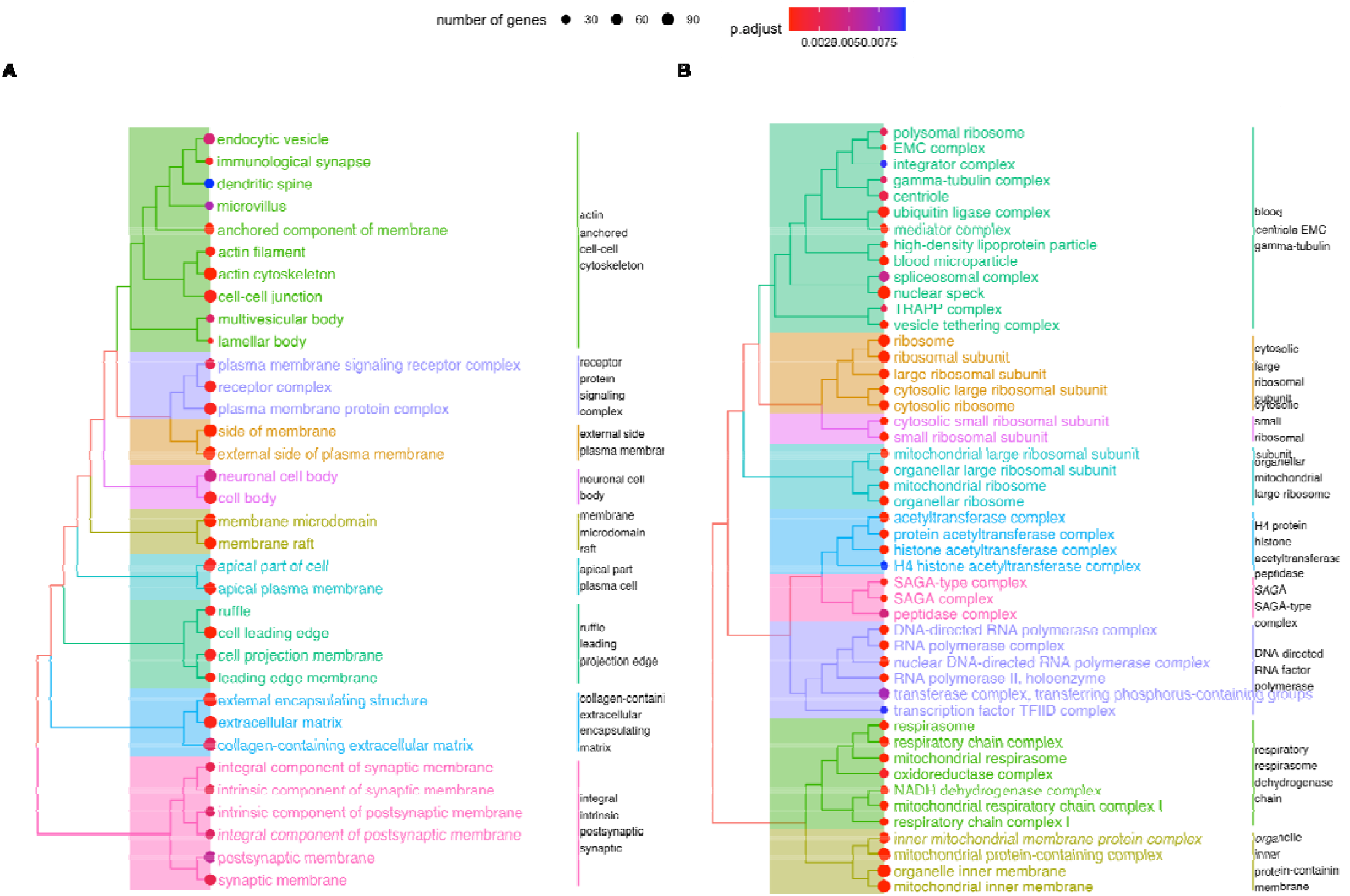
Pathway enrichment of proteins with good and poor predictability. **A.** Tree plots showing the clustering and relationships of gene ontology terms that are significantly enriched among proteins whose abundances are well predicted by their own transcripts (r ≥ 0.6). **B.** Tree plots of terms enriched among proteins whose abundances are poorly predicted by their own transcripts (r ≤ 0.3).

We next compared the genewise performances between the STRING feature elastic net model and the self transcript model to investigate the observed performance gain at a more granular level. Considering the 13,239 genes with protein-level prediction and with at least one STRING interactor, on average, each gene in the STRING model has a median of 273 [158–457] features, compared to 1 feature in the self transcript model. Incorporation of STRING features led to an average increase in test set correlation coefficients of 0.16 [IQR: 0.07–0.27] (**Supplementary Table S4**). There is a significant positive correlation between the number of features and prediction improvements (Pearson’s r 0.21, P: 8.2e–131) (**Supplemental Figure S3A**), a relationship which persisted when only genes with 10 or more interactors were included (**Supplemental Figure S3B**). A similar observation is seen in the CORUM feature set where the number of transcript features (i.e., CORUM complex interactors) of a particular protein is positively correlated with improvement in predictability of protein abundance (**Supplemental Figure S3C-D**). A functional enrichment analysis of proteins with strong improvements in prediction (difference in STRING vs. self-feature in test set predicted-actual correlation coefficient (Δr) ≥ 0.25) showed a strong enrichment in large multi-protein complexes including the ribosome, mitochondrial ribosome, RNA polymerase, and spliceosome (**Supplementary Table S5**) similar to the results from poor predictors. This corroborates that the non-correlation between transcripts and proteins of multi-protein complexes levels can be partially rescued by considering the information of interacting partner transcripts, which could participate in post-transcriptional regulations such as by being the stoichiometrically limiting transcript or crucial assembly parts, such that the transcript level of one subunit can regulate the protein level of other subunits.

Overall, these results are consistent with information about protein level residing both in the transcripts of the genes encoding the protein as well as the protein association partners. Moreover, taking into consideration the number of features in the STRING 800 and STRING 200 sets (median feature sizes of 10 [IQR: 2–48] and 266 [IQR: 149–451], respectively) compared to the size of the whole transcriptome feature set, the results indicate that a substantial portion of performance gains over the single-transcript feature set is already achievable from relatively few selected prior features.

### Proteins whose abundances are predicted by non-cognate transcripts are common

We next examined more closely the underlying causes behind the performance gains of protein predictions in the CORUM and STRING feature sets. To do so, we examined the genes whose protein prediction performance from transcripts increased substantially (Δr ≥ 0.25) after the incorporation of additional transcript features. In total, we observed 484 such proteins in the CORUM feature sets, and 3,272 proteins from the STRING data set, representing over 24% of all examined proteins. These numbers increase further when a less conservative Δr threshold of ≥ 0.15 is used (6,123 proteins in the STRING dataset, 946 in CORUM), altogether suggesting non-cognate transcript contributions to protein level are common at the proteome level. Upon inspecting the model feature coefficients (elastic nets) and feature importance (random forests), we observed that a considerable portion of these proteins with improved prediction are associated with (1) poor contribution from the self-transcript, and (2) a substantial contribution from primarily a few non-cognate (i.e., trans locus) transcripts coding for other proteins (see below). In other words, although model performance continued to increase with increased feature set sizes, the contributions of trans locus transcript features to overall prediction performance is unevenly distributed, and are therefore attributable to a few high contribution genes rather than a simple scaling with feature size. Notably, despite the STRING feature set being substantially larger than the CORUM feature set, substantial contributions from trans locus transcripts in the STRING feature set primarily involve transcripts encoding proteins that form part of the same CORUM complex as the protein of interest itself, suggesting that stable complex memberships play an outsized role in determining protein levels.

To illustrate the poor predictive power of cognate transcripts on some proteins, we next interrogated a subset of proteins where a non-self transcript has an outsized effect on the protein level of the proteins in the CORUM and STRING feature sets: Propionyl-CoA Carboxylase Subunit Beta (PCCB), C-X9-C Motif Containing 1 (CMC1), Proteasome Assembly Chaperone 2 (PSMG2), SMCR8-C9orf72 Complex Subunit (SMCR8), Mitochondrial Calcium Uptake 2 (MICU2), and Protein Phosphatase 3 Regulatory Subunit B, Alpha (PPP3R1). PCCB forms the propionyl-CoA carboxylase complex with two protein members, which breaks down certain amino acids in the cell (**Figure 3A**). PCCB showed poor prediction by their own self-transcripts (r:–0.008) but experienced high gains in test set correlation upon incorporating additional transcript features in the CORUM (r: 0.483) and STRING (r: 0.692) models. CMC1 is an assembly factor that forms an early intermediate of the cytochrome c oxidase complex in the mitochondrion with 14 documented protein members in CORUM (**Figure 3B**). Likewise, CMC1 showed only moderate prediction by their own self-transcripts (r: 0.274, respectively) but experienced high gains in test set correlation upon incorporating additional transcript features in the CORUM (r: 0.615) and STRING (r: 0.564) models.

**Figure 3.**
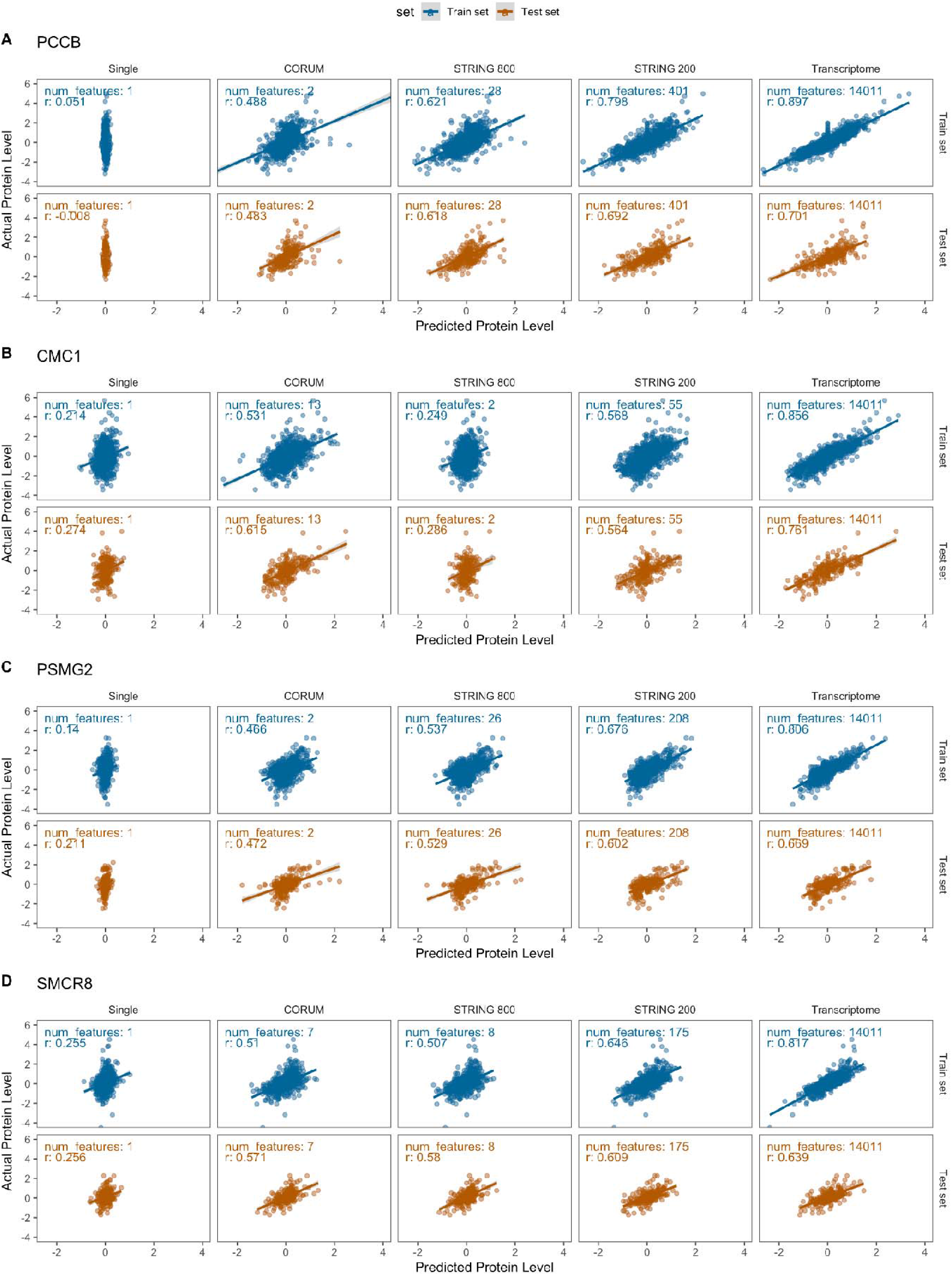
Proteins with improved predicted levels after inclusion of additional transcript features. Four proteins with substantial predictability from transcriptome data upon the inclusion of additional features are shown: **A.** PCCB, **B.** CMC1, **C.** PSMG2, **D.** SMCR8. For each protein, the transcript-trained prediction of protein level is plotted on the x axis and the actual protein level is plotted on the y axis. The lack of variance in predicted protein levels from the self-transcript model is due to the regularization of the elastic net model, and corresponds to a lack of correlation between PCCB mRNA and protein (see Figure 4). Blue: train set, brown: test set. Columns denote the transcript feature set used to train the model. The number of features used to train the model in each feature set is shown inside each plot. r: Correlation coefficient.

We then considered the transcriptomics and proteomics data distributions of the features that had highest coefficients in the elastic net model from the CORUM feature set for PCCB and CMC1. For PCCB, most protein-level predictions are recovered when the feature set includes PCCA, which together with PCCB forms the stable propionyl-CoA carboxylase enzyme that consists of six copies of PCCB and six copies of PCCA each. There is a remarkably strong correlation with its propionyl-CoA carboxylase complex interacting partner PCCA at the protein level (r: 0.957), but this correlation is almost entirely absent at the transcript level (r: −0.059). This is partially explainable by the observation that PCCA transcript has a higher level of variability and is also strongly correlated with the PCCB protein level. The total protein level of PCCB is therefore primarily driven by PCCA rather than PCCB transcripts (**Figure 4A**).

**Figure 4:**
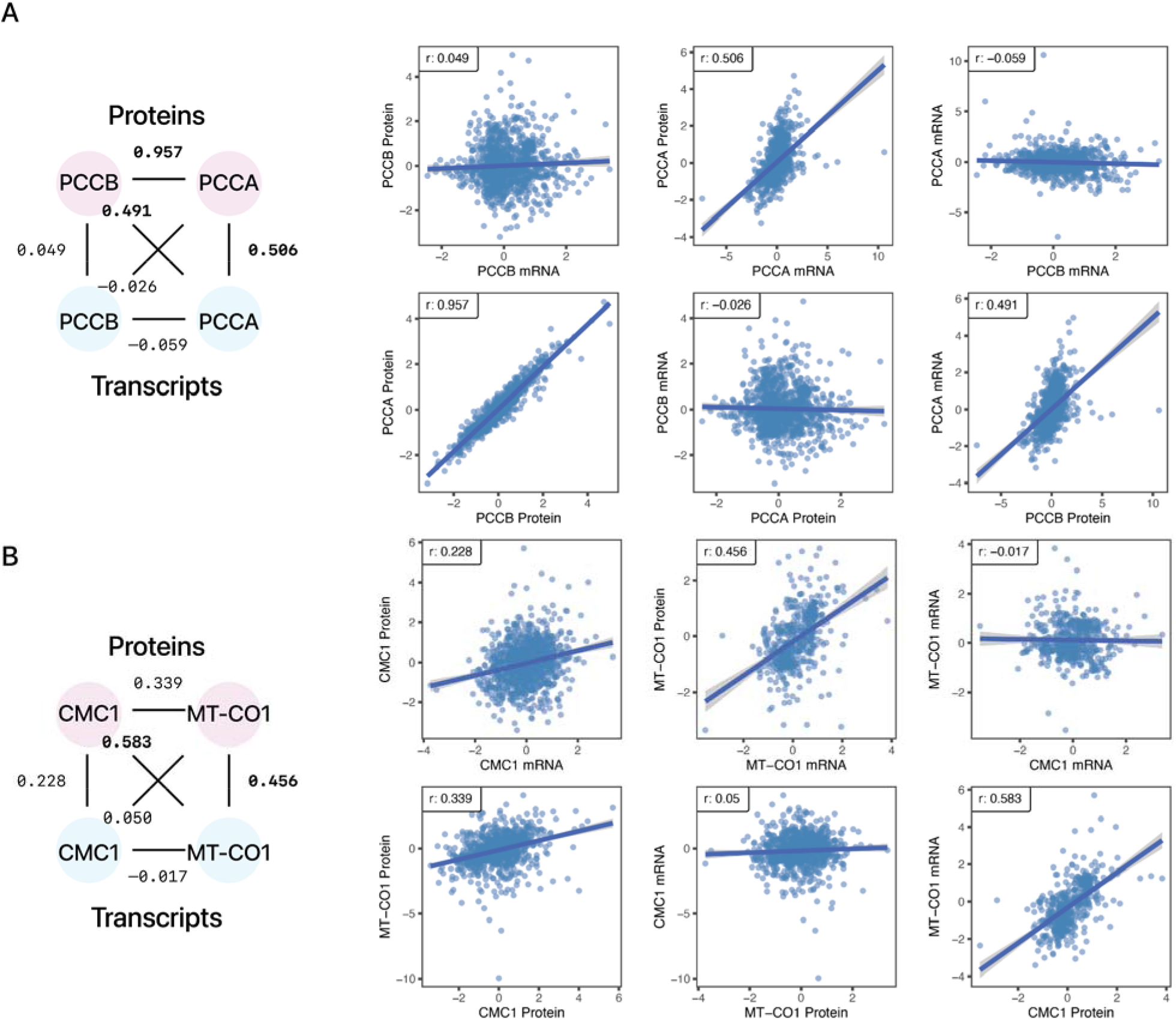
mRNA-Protein correlations of PCCB and CMC1 with functionally associated proteins. Two examples of proteins whose abundance is better explained by another transcript are shown. **A.** PCCB protein level is predicted by PCCA transcript but not its own transcript. **B.** CMC1 protein level is explained by MT-CO1 transcript level but not its own transcript. Substantial correlations across transcripts and proteins (≥ 0.4) are bolded.

In the case of CMC1, we found that MT-CO1 has the highest contribution to CMC1 protein level (**Figure 4B**). MT-CO1 is the mitochondrial genome encoded subunit of complex IV that is thought to act as the nascent scaffold around which the complex is assembled. CMC1 binds to MT-CO1 during the formation of the early module known as MITRAC and is subsequently released during assembly. ^43,44^ CMC1 transcript and protein levels are moderately correlated with one another (r: 0.228). Again, CMC1 and MT-CO1 show poor co-expression at the transcript (r: −0.017) level and an improved correlation at the protein (r: 0.339) level, whereas MT-CO1 transcript is robustly correlated with the CMC1 protein level (r: 0.583), which suggests the possibility that proteogenomic co-expression may reveal additional functionally related protein pairs. Closer inspection suggests the possibility of a non-linear relationship between MT-CO1 transcript and CMC1 protein where upon reaching a plateau, further increases in MT-CO1 transcript does not correlate to further increase in CMC1 protein levels, suggesting other transcripts may also contribute to CMC1 protein levels. Consistent with this, other complex IV subunits also have non-negligible coefficients (in the elastic nets) or feature importance (in the random forests) in the CMC1 model. When the feature set expanded to include all STRING proteins, MT-CO1 remained a high contributor in the random forest but not the elastic net model, suggesting the elastic nets may be more susceptible to collinear features than the random forests.

Two other examples of small stable complexes with interdependent protein and mRNA correlation are shown in **Figure 5.** Proteasome Assembly Chaperone 2 (PSMG2) forms the heterodimer PAC1-PAC2 complex (2 protein members in CORUM) along with Proteasome Assembly Chaperone 1 (PSMG1), which binds with 20S proteasome precursors and acts as a scaffold to promote proteasome ring assembly while preventing aberrant dimerization of 20S proteasome α rings ^45^. PSMG2 and PSMG1 are co-expressed strongly at the protein level (r: 0.713) but only moderately at the transcript level (0.222) (**Figure 5A**). PSMG2 protein has a higher correlation with PSMG1 transcript (r: 0.472) than its own (r: 0.158). From GTEx v8 data, PSMG2 is expressed more highly than PSMG1 across tissues (TPM ~40 vs. 15) ^46^, which suggests PSMG1 may act as a limiting factor in heterodimer formation. PSMG1 knockout in mice has been shown to decrease PSMG2 at the protein level, consistent with our finding that PSMG1 transcript levels influence PSMG2 protein abundance ^47^.

**Figure 5:**
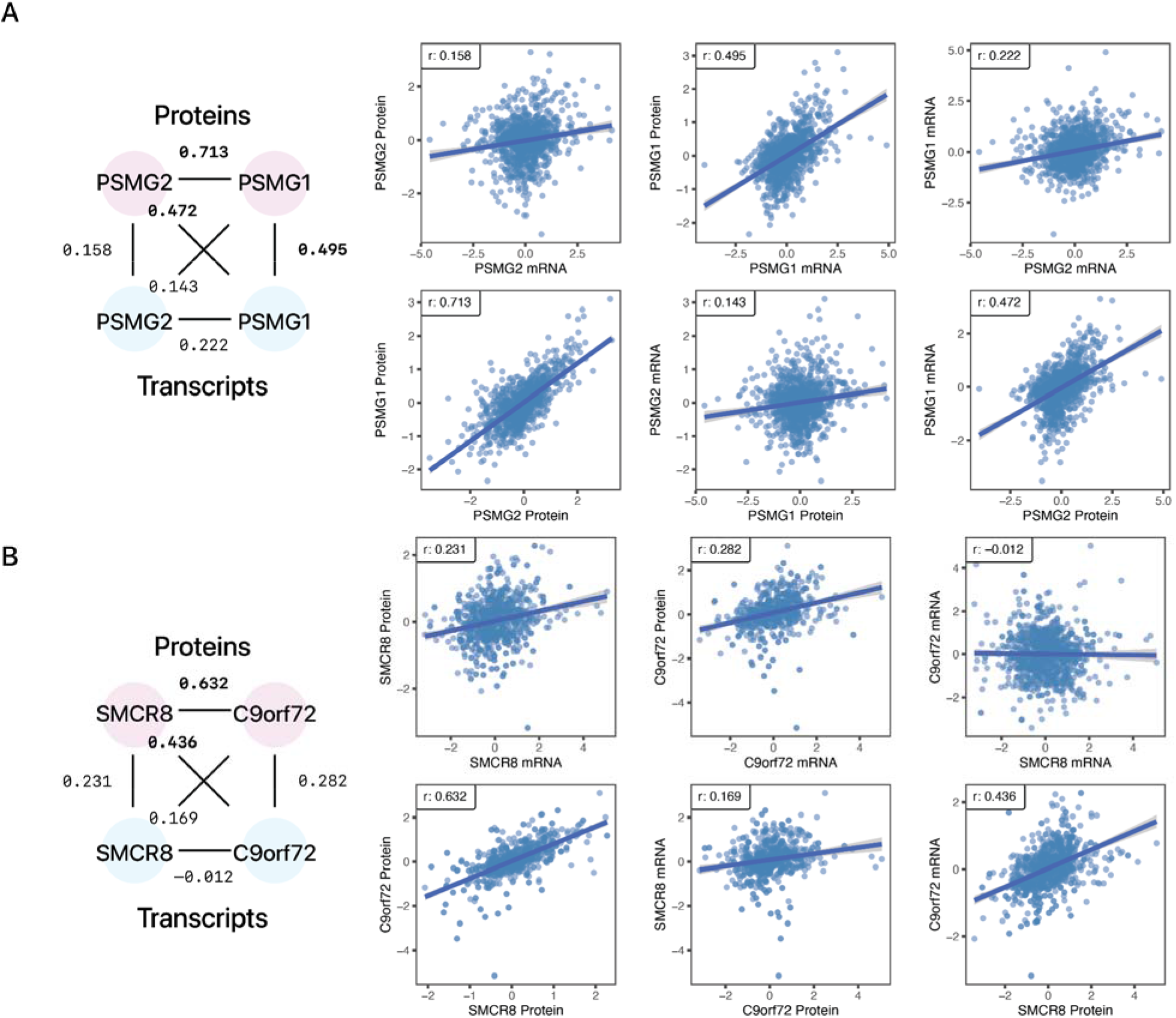
mRNA-Protein correlations of PSMG2 and SMCR8 with functionally associated proteins. Two examples of proteins whose abundance is better explained by another transcript are shown. **A.** PSMG2 protein level is predicted by PSMG1 transcript but not its own transcript. **B.** SMCR8 protein level is explained by C9orf72 transcript level but not its own transcript. Substantial correlations across transcripts and proteins (≥ 0.4) are bolded.

In another example, Smith-Magenis Syndrome Chromosomal Region Candidate Gene 8 (SMCR8) forms the heterotrimer C9orf72-SMCR8-WDR41 complex (3 protein members in CORUM) together with the C9orf72-SMCR8 Complex Subunit (C9orf72) and WD Repeat Domain 41 (WDR41) ^48^. C9orf72-SMCR8 is known to dimerize prior to binding with WDR41 to form the heterotrimer. The C9orf72-SMCR8 complex modulates autophagy, likely by modulating the maturation of autophagosomes ^49^. Mutations in the genes encoding the C9orf72-SMCR8 Complex are broadly implicated in diseases including amyotrophic lateral sclerosis and frontotemporal dementia. As in the other cases, we observed a co-expression of SMCR8 and C9orf72 at the protein level (0.632) but not the transcript level (–0.012), and SMCR8 protein levels correlate more strongly with the C9orf72 transcript (r: 0.436) than the SMCR8 transcript (r:0.231) (**Figure 5B**). A previous study suggested C9orf72 may stabilize excess SMCR8 ^50^, providing one possible explanation for the mechanism behind the observations here.

**Supplementary Figure S5** shows two more examples concerning small complexes. The mitochondrial calcium uniporter complex (MCU complex) contains five protein members in CORUM (**Supplementary Figure S5A**). Regulatory subunits of the complex, including calcium uptake protein 2, mitochondrial (MICU2) sense calcium levels, regulating the uptake of calcium performed by the mitochondrial calcium uniporter (MCU) ^51^. Heterologous overexpression of FLAG-tagged MCU has been shown to increase expression of MICU2 in culture ^52^. This is interesting, given the observation here that at an endogenous level of MCU expression, MICU2 protein is more strongly correlated to the MCU transcript (r: 0.488) than its own (r: 0.058) (**Supplementary Figure S5A**).

The final example highlighted here comes from the heterodimer calcineurin, which is made up of a catalytic subunit (calcineurin subunit A), with 3 isoforms encoded by 3 separate genes (PPP3CA, PPP3CB and PPP3CC), [CJ2] and a Ca^2+^ binding regulatory subunit (calcineurin subunit B), with 2 isoforms also encoded by 2 separate genes (PPP3R1 and PPP3R2) ^53^. Calcineurin is a serine/threonine phosphatase which modulates many calcium dependent signaling pathways, and is the target of immunosuppressant drugs cyclosporin A and FK506 ^54^. In the analysis here, calcineurin subunit B type 1 (PPP3R1) protein correlated poorly with PPP3R1 transcript levels (r: 0.205), but strongly correlates (r: 0.819) with the protein phosphatase 3 catalytic subunit alpha (PPP3CA) transcript –– an isoform of calcineurin subunit A (**Supplementary Figure S5B**). Together, these cases illustrate a general observation –– in cases where the protein abundance of a gene is more highly correlated to the transcript level of a binding partner than to its cognate transcript, there appears to be stronger co-expression between the gene and its abundance driver at the protein level but not at the transcript level.

To corroborate these observations, we used the Boruta algorithm against the random forest CORUM models and confirmed each of the principal non-cognate transcript features is retained. We further considered Shapley values as an explanation of random forest and gradient boosting models, and likewise found that the interacting protein transcripts formed the top contributors in explaining the abundance of the protein of interest in each of the six cases above (**Supplementary Figure S6**). In some examined cases including PSMG2, the transcript coding for the protein of action is more abundant than that of its regulating interacting partner, which is compatible with a scenario where simple stoichiometric constraints control the protein level of the supernumerary subunit. It is however less clear whether this scenario applies to other examined cases including CMC1 and MT-CO1, where the MT-CO1 transcript is much more abundant than CMC1 owing to the multiple copies of mitochondrial genome in the tissue. Hence, how MT-CO1 regulates CMC1 protein level is not explainable directly from transcript stoichiometry alone and awaits further mechanistic studies. Finally, we also examined feature importance in a single data set of a recent CPTAC study that showed the highest performance in the single feature model (LSCC) (**Supplementary Figure S7**). Among the six highlighted proteins (PCCB, CMC1, PSMG2, SMCR8, MICU2, PPP3R1), five were best predicted by a non-self transcript, and five of the top trans locus predictors were conserved from the combined data set analysis with the exception of MT-CO1, which was not among the model features due to the number of shared observations required. Hence the observed trans locus predictors are conserved in a single data set model and are unlikely to be due to heterogeneity across data sets.

**Table 1** summarizes the top 50 genes whose protein abundance becomes substantially more predictable upon the inclusion of CORUM features, along with their top protein level contributor transcripts. Notably, a number of these genes are found in multiple gene sets within the MSigDB C2 CGP gene set collection that are commonly used for functional enrichment analysis ^55^, whereas others are found to be significantly associated with diseases in the literature from bibliometric analysis ^56,57^, hence these genes are common and important to multiple areas of biomedical inquiries. We suggest the possibility that the transcripts of these genes may not present accurate proxy variables for their proteins should be considered in linking transcript level changes to downstream cellular physiology.

**Table 1:**
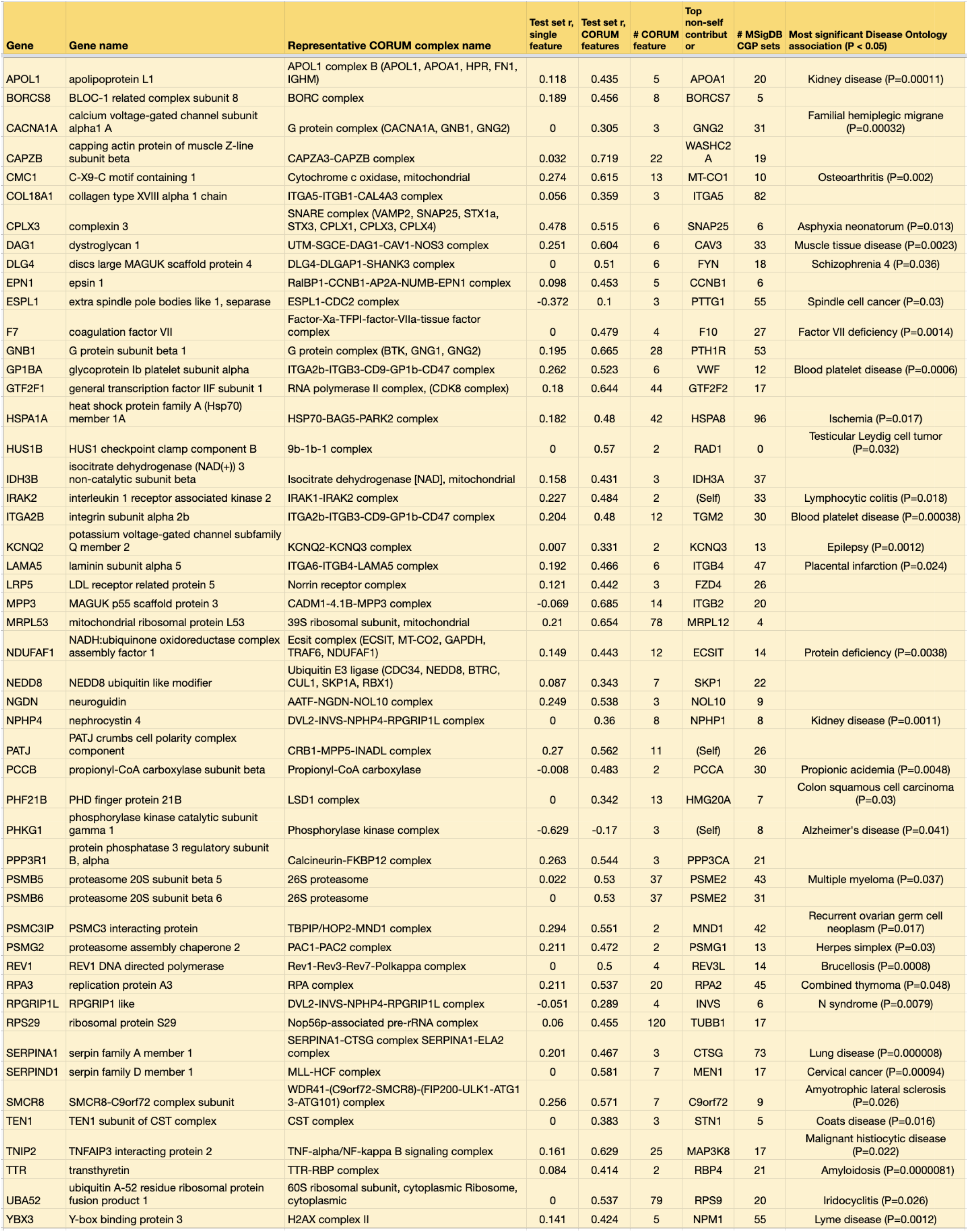
Top 50 proteins whose abundance is under substantial influence from non-cognate transcripts. Columns 1 and 2: gene names. Columns 3 show the representative CORUM complex the gene of interest belongs to. Columns 4 and 5 denote the increase in prediction performance between elastic net single feature (self transcript) vs. CORUM feature sets. Column 6 shows the number of transcripts used to predict the protein level of the gene of interest in the CORUM feature set. Column 7 shows the top trans-locus contributor to the protein level of the gene of interest, ranked by absolute coefficients in the elastic net model. Proteins whose own transcripts are the top predictors are marked with (self). Column 8 denotes the number of MSigDB C2 CGP (chemical and genetic perturbation) gene sets in which the gene appears. Column 9 denotes the top significantly associated Disease Ontology term with the gene of interest in the literature.

Taken together, these results exemplify widespread interdependency of protein levels on trans locus transcripts, involving not only large megadalton protein complexes but many small complexes involved in diverse biological processes in the cell. The CORUM feature set considered here alone contains over 2,700 human complexes derived from CORUM release 3.0 mappable to 3,689 proteins with distinct gene names, with a median of 3 proteins per complex (**Supplementary Figure S8**). Thus, small protein complexes are widespread in the proteome and a large fraction of the proteome could be placed under complex post-transcriptional control that decouples protein levels from mRNA levels. Moreover, half of the proteins in the CORUM annotations belong to 2 or more distinct complexes (including intermediates and subcomplexes) which suggest further opportunities for more complex patterns of interdependent protein and mRNA levels.

### Network representation of the interdependencies of protein level regulations

We next examined whether the interdependency of proteins and trans locus transcripts may be used to infer potential novel regulatory drivers. To do so, we constructed graphical models to visualize the connections between each protein and the transcriptome features that contribute to its predicted level, followed by topological analysis to find hub nodes and extract network patterns. To limit scope, we generated graphs to proteins whose prediction improved in the CPTAC_8 data set following the inclusion of the CORUM or STRING features (Δr ≥ 0.25) A directed graph is then generated using a list of edges that connect feature variables (transcripts) to target variables (proteins), with the edge weights calculated using a function of the elastic net coefficients and random forest feature importance of the models for both the CORUM (**Supplementary Table S6**) and the STRING (**Supplementary Table S7**) feature sets. The resulting network is partitioned into connected components of transcript-protein relationships. The subgraph that corresponds to the relationship between PCCA and PCCB as discussed above shows the PCCB node contributing (as transcript) to its own protein level weakly, and the PCCA node contributing strongly to the PCCB node (as protein) (**Figure 6A**). The relationship of CMC1 with other cytochrome c oxidase complex subunits, as described above, is represented in a wheel and spokes pattern where multiple transcripts (orange nodes) contribute either positively or negatively to the CMC1 protein level, with MT-CO1 having the strongest positive contributions (**Figure 6B**). Another member of the MITRAC module COA3 to which CMC1 binds likewise exerts a positive influence on the protein level of CMC1, consistent with the possibility that interdependent protein and transcript relationships may reflect information related to protein complex assembly sequences. Larger subgraphs represent instances where two or more protein target nodes are connected by shared transcript source nodes. Due to the method by which the graphs are generated, protein hubs (with high in-degrees) are more prevalent, but transcripts with high outflow activities are also seen that connect two or more protein nodes. A subgraph in **Figure 6c** predicts that the transcript levels of SH3KBP1 and CD2AP both contribute to the predicted protein levels of multiple proteins involved in the RICH1/AMOT polarity complex and the PI4K2A-WASH complex, and the protein level of WASHC1 is influenced by genes in the CCC-Wash (WASH1, FAM21C) complex and the PI4K2A-WASH complex (**Figure 6C**).

**Figure 6:**
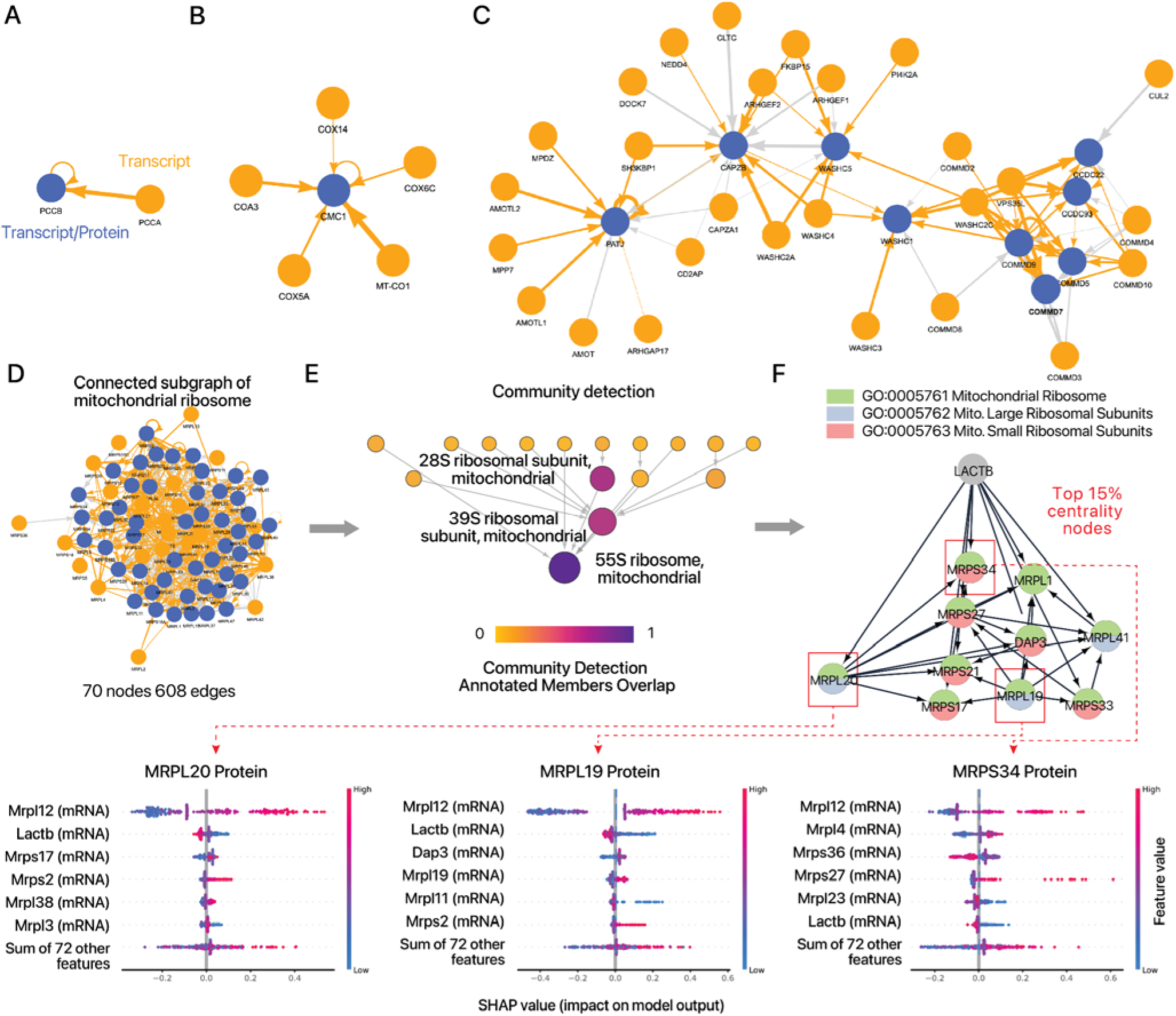
Directed graphs of protein and transcript interrelationships identify candidate regulatory genes. **A-C.** Examples of directed graphs constructed from genome-wide relationships of transcript-predicted proteins, containing members of **A.** the propionyl-CoA carboxylase complex; **B.** the cytochrome c oxidase, mitochondrial complex; **C.** the PI4K2A-WASH complex, the RICH1/AMOT polarity complex, and others. In each subgraph, orange nodes have outflow edges only (i.e., they are contributing transcripts in the prediction models). Blue nodes are nodes that are connected to other nodes via at least one inflow edge (i.e., they represent proteins, and optionally also transcripts if they also have outward edges). Orange edges represent positive coefficients of the transcripts to the target proteins in the elastic net models; gray edges represent negative coefficients. All edges are directed from transcript to protein, and the widths of the edges are scaled by the weight. **D.** A highly connected subgraph of mitochondrial ribosome subunits containing 73 nodes and 834 edges. **E.** Persistent community detection and network representation of preferential node connections, showing a hierarchical relationship between the 28S and 39S subcomplex with the assembled 55S mitochondrial ribosome. **F.** Network representation of hub nodes defined as 15% of nodes ranked by betweenness centrality, which predicts a potential role of LACTB as a critical hub that lies upstream of multiple large and small mitochondrial ribosomal protein subunits. Node colors represent the pie chart diagram of the corresponding GO biological process described in the table. SHAP values of three proteins (MRPL20, MRPL19, MRPS34) are highlighted showing top model contributors.

We highlight an instance where mRNA-protein relationships may be examined to generate new hypotheses. The mitochondrial ribosome is represented in a highly connected subgraph where the abundances of a majority of complex subunit proteins are each contingent upon multiple transcripts (**Figure 6D**). Using topological analysis based on the hierarchical community detection framework (HiDeF) method ^33^, we showed that this subgraph preserves the expected hierarchical relationship between the 28S small and 39S large subunits in forming the 55S mitochondrial ribosome, suggesting the preferential connections within the proteogenomic graphs broadly recapitulate subcomplex assembly (**Figure 6E**). We then extracted the hub nodes from the subgraph using the betweenness centrality algorithm, which revealed mitochondrial serine beta-lactamase-like protein (LACTB) to be an upstream gene that is highly connected within the subgraph and exerts a regulatory effect on both small and large mitochondrial ribosome subunit components. Feature importance analysis further suggests the *LACTB* transcript to have a negative impact on multiple ribosomal mitochondrial proteins (**Figure 6F**). Of note, *LACTB* was previously identified as a new subunit (MRP-L56) of the mitochondrial ribosomal large subunit isolated from sucrose gradients ^58^. Subsequent work however has established the LACTB protein as a filament-forming protein that is localized to the intermembrane space instead away from the mitochondrial ribosomes ^59^ that possesses in vitro protease activity ^60^ and that acts as a tumor suppressor by maintaining post-mitotic differentiation states ^60^. The physiological function and antiproliferative mechanism of LACTB are incompletely understood, although induced LACTB expression in cancer cells was found to affect mitochondrial phospholipid metabolism ^60^. The analysis here therefore raises the possibility that *LACTB* may also affect oxidative phosphorylation in differentiated cells by regulating mitochondrial ribosome biogenesis and protein synthesis, which may be validated experimentally.

A similar hub node analysis from the largest subgraph from the STRING feature set (**Supplemental Table S8**) showed chromogranin A and B (CHGA and CHGB) as inward hubs (i.e., proteins associated with many transcripts) associated with secreted peptides including CST3, SERPINC1, MFGE8, SERPIND1, and others (**Supplementary Figure S9**), which is consistent with the known roles of these acidic glycoproteins as the primary constituents of secretory granules that can regulate their rate of their formation^52^. Taken together, the topological analysis shown here suggests mRNA-protein networks may be useful for generating new hypothesis on regulatory drivers from large proteogenomics data.

## Discussion

### Widespread contributions of trans locus transcripts to protein level

Recent work has applied various machine learning models to the computational task of predicting across-sample protein levels using transcriptomics data, but investigation into the biological factors that uncouple transcriptome and proteome data have remained limited, noting only that predictability may differ across broad functional categories, e.g., essential genes or metabolic genes may be less predictable. The results here suggest that the transcript levels of interacting partners had an outsized contribution to the abundance of proteins of interest. The notion that protein-protein interaction can influence protein abundance post-transcriptionally is not new, as it is known that supernumerary subunits of protein complexes can be removed through protein degradation. However, details of genes that show especially poor mRNA and protein abundance correlations and the identities of their protein regulators have remained scarce. The current study provides new evidence from protein prediction models that protein abundance by other transcripts is very common in the proteome, nominating specific interactions involving not only large megadalton sized multi-protein complexes as previously observed, but also smaller stable complexes (e.g., propionyl-CoA carboxylase with two subunits) in the CORUM feature set and potentially more transient interactions documented in the STRING feature set. The common presence of small stable complexes in the proteome greatly expands the repertoire of proteins for which due considerations should be given when directly interpreting transcript level data as representing protein level information. Moreover, it has been suggested that promiscuous protein-protein interactions without established biological function may be a common occurrence during the co-evolution of functional protein-protein interactions ^62^. The occurrence of such stable but non-functional interacting pairs could further increase the scope of trans locus protein regulation which would have implications on the predictability of protein levels from mRNA levels.

The imperfect correlation between protein and mRNA points to orthogonal information that exists between transcript and protein regulations, which underpins the untapped potential for further multi-omics integration to derive new insights ^63^. The use of protein correlations to find causal insights for post-transcriptional regulations has previously been explored in other contexts, which looked for anti-correlation between E3 ubiquitin ligases with known proteins or ubiquitination sites of interest, which may control their protein level by virtue of post-translational degradation ^18,24^. Extending this intuition towards multi-omics correlation, we used directed graphs generated from the model coefficients and feature importance of trans locus transcripts to represent the interaction patterns between the transcripts and proteins of interest. This in turn nominated a number of hub proteins whose abundance is contributed by multiple transcripts and hub transcripts that regulate multiple proteins, and enabled community detection analysis to find the relationships between biological processes in protein and mRNA correlation networks. As more data continue to become available, we foresee that graphical models will be useful for finding more trans locus regulations of protein levels, such as those that represent known assembly sequential steps or post-transcriptional regulators. These graphical models may be used to generate testable hypotheses or find utility in predicting experimental outcome, barring confounders or reverse causality effects. For instance, in the case of the actin related protein 2/3 complex, one might predict that within a certain concentration range an overexpression of *ACTR3* and *ARPC4* will be more effective in modulating ARPC3 protein levels than augmenting the expression of *ARPC3* itself, which is readily testable by experimentation.

### Limitations of the study

Significant differences in protein regulations likely occur in different cell and tissue types. The predictive models here are trained using publicly available CPTAC data from 8 cancer types, which contain transcriptome and proteome data from both tumor and normal adjacent tissue samples. In prior work, we found general concordance between protein and mRNA correlation in CPTAC samples vs. GTEx tissue proteomics data ^15^. However, other studies that compared protein and mRNA correlation in tumors and normal adjacent tissues have found higher inter-sample correlation in tumor samples ^21,22^, which may be attributable to the increased translation rates in cancer. Hence, additional discordant cases between proteins and mRNA likely remain to be discovered that are omitted here. Sample difference may also explain the observation from GTEx proteomics that secreted proteins are associated with protein and mRNA discordance ^7^ as one might intuitively assume but which is not apparent in the analysis here. The cases used for training the model are not labeled by their cancer type or their tumor vs. normal adjacent tissue designation, which would likely have availed overall predictive performance.

We performed the analysis using expression data from TCGA/CPTAC RNA sequencing experiments through the cptac Python API, which retrieves the final data tables from the flagship CPTAC papers of each individual cancer type ^25^. Although each TCGA/CPTAC cancer subtype project follows an overall consistent experimental design and data acquisition strategy, minute differences exist in the processing pipelines used to analyze the RNA sequencing data (e.g., STAR vs. Bowtie2) and gene expression measure (e.g., RPKM vs. FPKM) which could bias gene expression values across cancer types. Likewise, there are subtle differences in the protein expression data in each cancer type (e.g., search engine, isotope tags). As our goal here is not to compare trends across cancer types, we have taken inspiration from the CPTAC Dream Challenge submissions that improved overall predictions by borrowing information across cancer types. However, it is also known that different data sets may present different variability. Future work may employ uniformed processed data to improve performance and reduce bias, such as data from the TCGA PanCanAtlas which processed all TCGA cancer RNA sequencing data uniformly ^64^, or compare batch correction strategies. To our knowledge, uniformly processed protein data are not yet available at the time of writing. Other machine learning and deep learning algorithms can likely further boost protein level predictions. We have limited our scope here to comparisons of different feature sets and previously employed algorithms, and to result interpretation.

Finally, pitfalls must be considered when attempting to interpret predictive models in search of mechanistic insights. Feature importance in the predictive models represent correlation rather than causality, and hence interpretations can be confounded by independent confounders, multicollinearity, and reverse causality. In simple cases such as PCCB–PCCA, given the emphasis on prior feature selections that prioritize known protein-protein interaction partners, the logical interpretation would be to assume the hierarchical nature of gene regulations where transcript levels are more likely to affect protein abundance than the opposite, but this assumption becomes more tenuous as feature sets expand and the number of associations increase, and in cases such as transcription factor proteins whose abundance can affect transcript levels across samples. Future work in this area may employ more sophisticated causal inference methods to identify regulatory modalities.

## Conclusion

In summary, this study compared predictive models of protein levels using different transcript feature sets, and provided biological interpretations of the results by highlighting trans locus transcripts with substantial contributions to protein levels. Although the concept of proteins being modulated by their interacting partners is not new, to our knowledge the prevalence of these regulations have not been examined at a proteome wide level and using recently available proteogenomics data sets. The analysis here therefore reveals new details into the gene identity and modality of trans regulation of protein levels, and gives support to further development of prior transcript feature selection strategies to optimize protein prediction tasks. The results show that the transcript levels of protein-protein interaction partners can broadly influence protein abundance in a tissue, which has implications on the interpretations of transcriptomics data and on understanding the architecture of proteome composition regulations. With further refinement of feature selection and feature engineering methods and the availability of large data sets, we foresee that similar approaches to those shown here will provide valuable new insights into post-transcriptional mechanisms of protein regulations.

## Author Contributions

EL conceptualized the project. HS and EL wrote software code. HS, MJL, and EL performed data analysis. HS, MJL, JC, RC, MPL, and EL drafted the manuscript. MPL and EL revised the manuscript.

## Acknowledgments

This work was supported in part by NIH Office of the Director award R03-OD032666; NIH/NHLBI award R00-HL144829 to EL; and NIH/NHLBI awards R00-HL127302, R01-HL141278, and the Consortium for Fibrosis Research & Translation funds at the University of Colorado to ML.

## Supplementary Figures

**Supplementary Figure S1:**
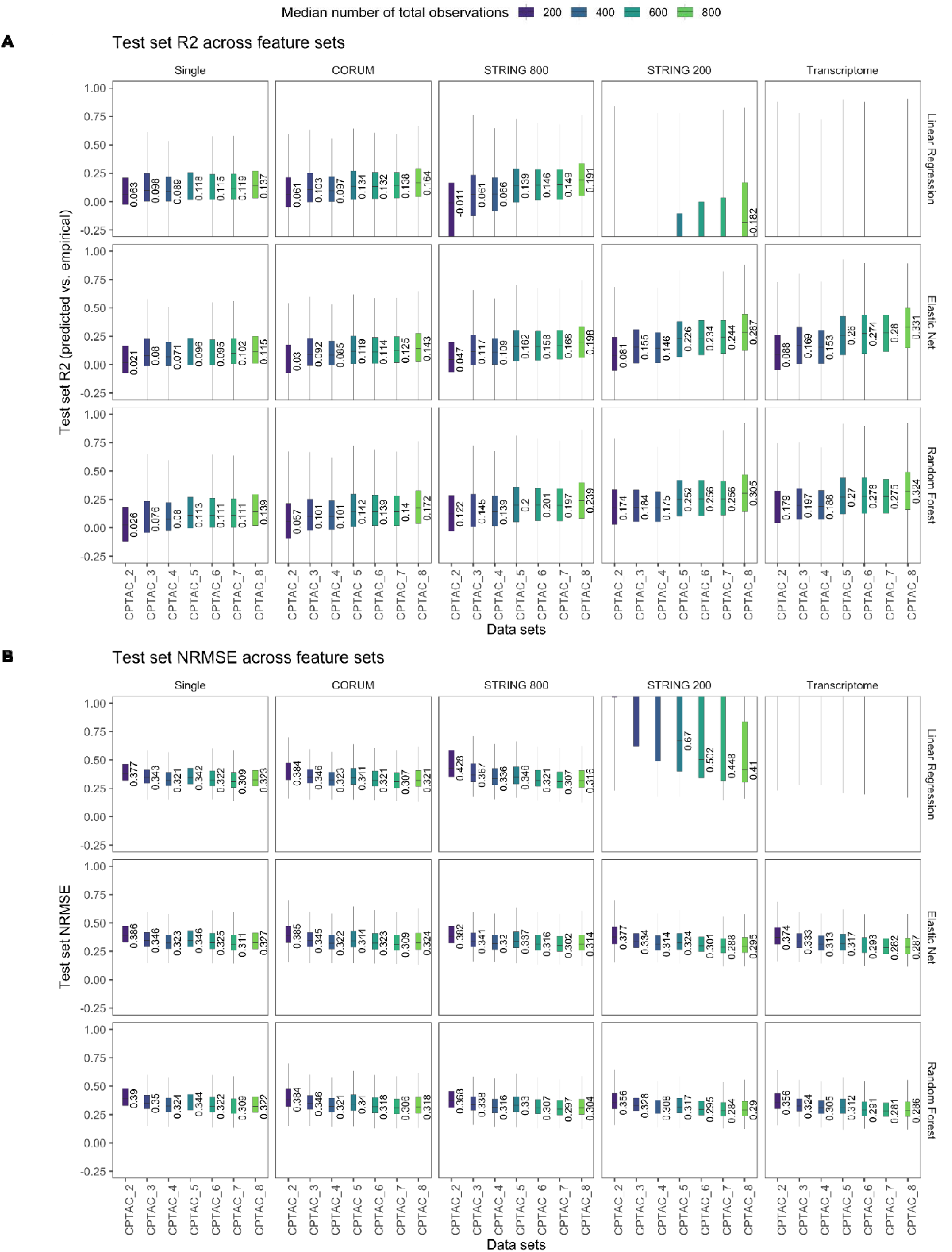
Model performance measured by additional metrics. **A.** Box plots of test set normalized root mean square error (NRMSE) between the transcript-predicted and actual protein level for each protein are shown across five feature sets (column: single/self transcript, CORUM interactors, STRING high-confidence associated proteins; STRING low-confidence associated proteins, and all transcripts) and three algorithms (multiple linear regression, elastic net, and random forest). In each plot, x axis denotes the number of CPTAC data set used to train the models box: interquartile range; whiskers: +/− 1.5 IQR. **B.** As above, but for test set goodness-of-fit (R^2^).

**Supplementary Figure S2:**
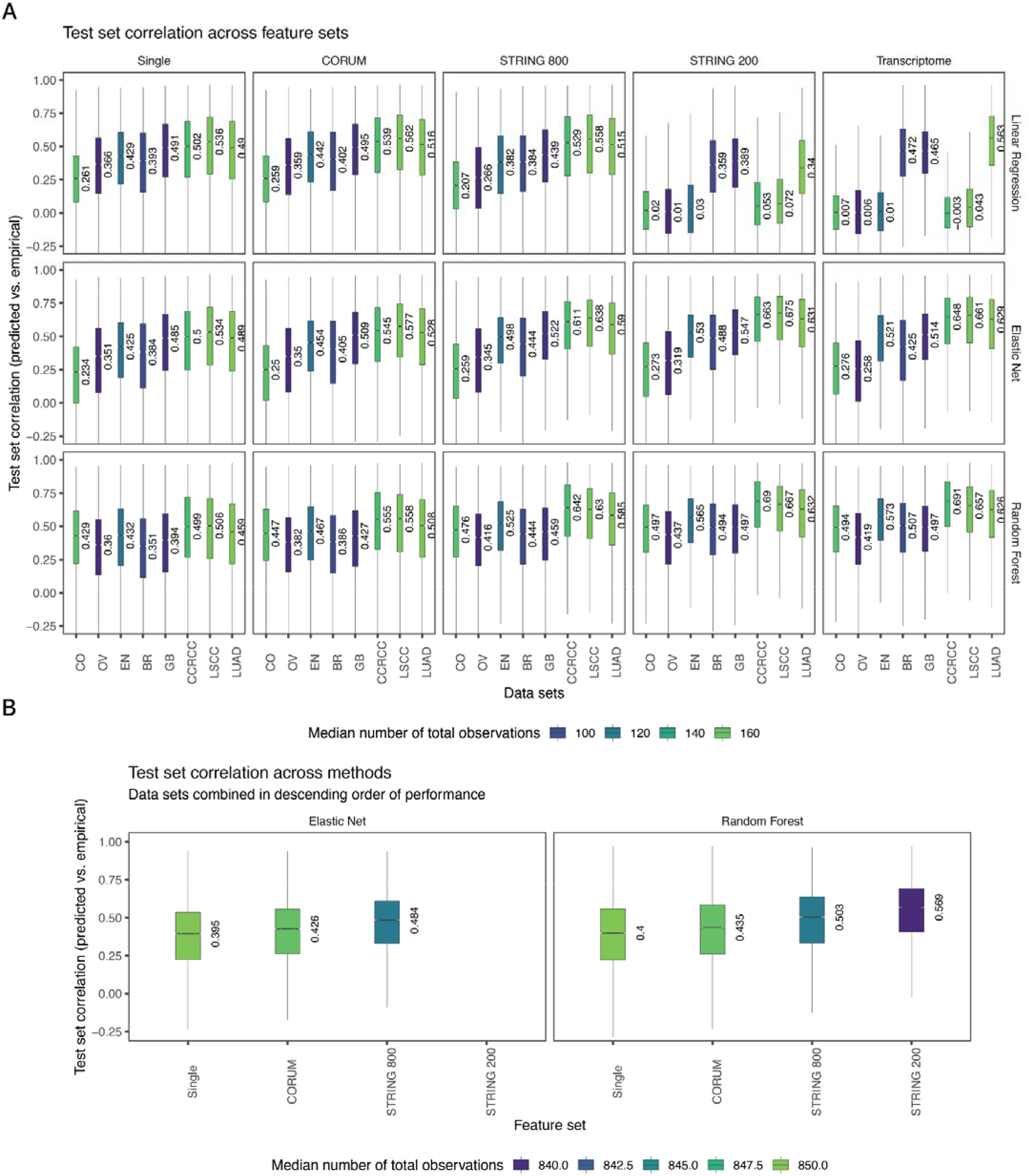
**A.** Model performance in single cancer data sets. Box plots of test set correlation coefficients (r) between the transcript-predicted and actual protein level for each protein are shown across five feature sets (column: single/self transcript, CORUM interactors, STRING high-confidence associated proteins; STRING low-confidence associated proteins, and all transcripts) and three algorithms (multiple linear regression, elastic net, and random forest). In each plot, x axis denotes the CPTAC cancer type study used to train the models; box: interquartile range; whiskers: +/− 1.5 IQR. **B.** Model performance when the 8 data sets were combined in the order of decreasing single data set performance.

**Supplementary Figure S3.**
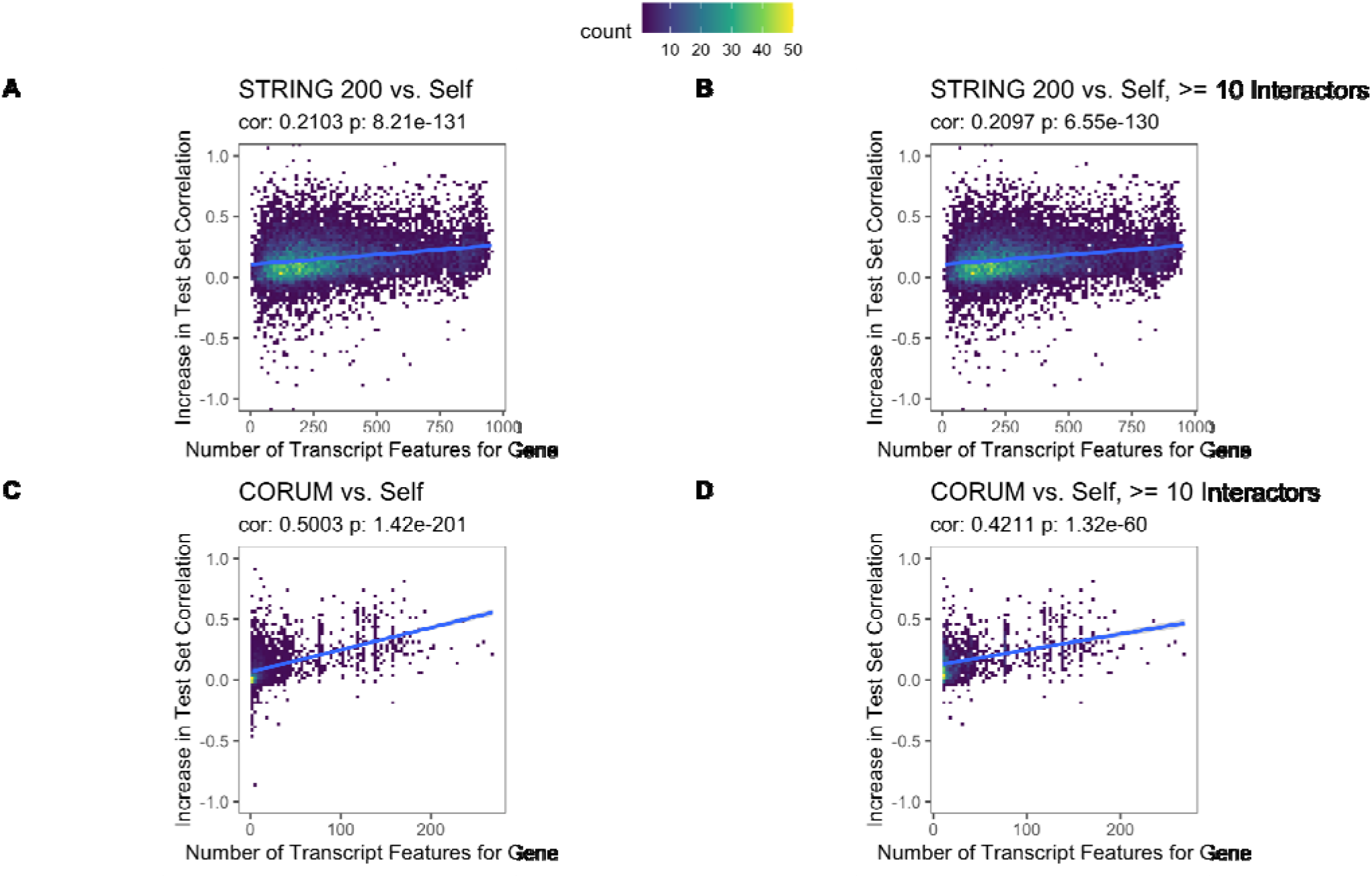
Improvements to protein prediction from the incorporation of additional transcript features. **A–B.** Scatterplot showing a significant linear relationship between the number of low-confidence interactors a protein has as annotated in STRING vs. the increase in test set correlation between predicted vs. actual protein levels in the Elastic Net models over the self-transcript feature set (correlation test p: 8.2e–131 for all proteins; 6.5e–130 for proteins with 10 or more interactors). **C–D.** As above, but for the CORUM feature set.

**Supplementary Figure S4:**
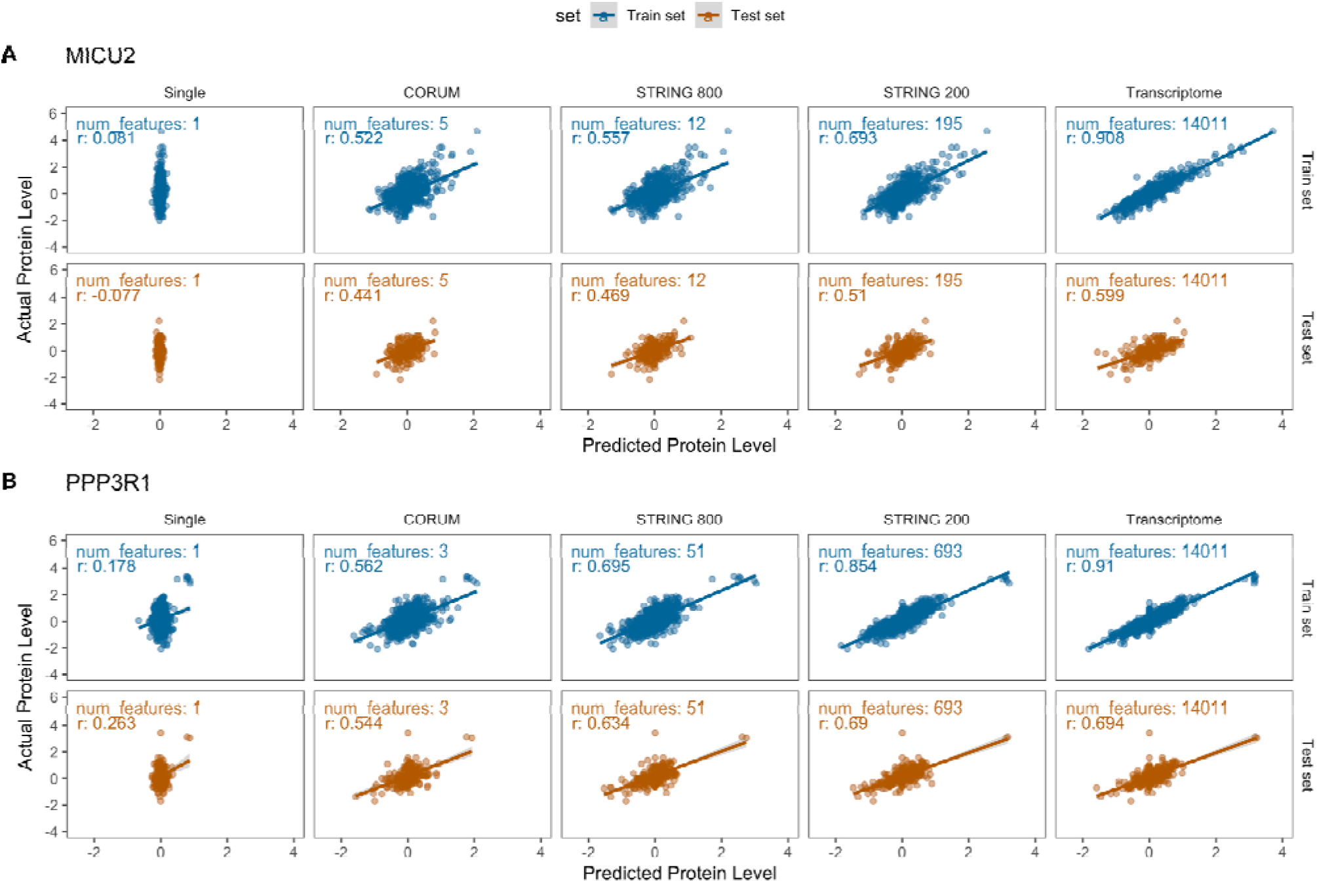
Additional proteins with improved predicted levels after inclusion of additional transcript features. Two additional highlighted proteins with substantial predictability from transcriptome data upon the inclusion of additional features are shown: **A.** MICU2, **B.** PPP3R1. For each protein, the transcript-trained prediction of protein level is plotted on the x axis and the actual protein level is plotted on the y axis. Blue: train set, brown: test set. Columns denote the transcript feature set used to train the model. The number of features used to train the model in each feature set is shown inside each plot. r: Correlation coefficient.

**Supplementary Figure S5:**
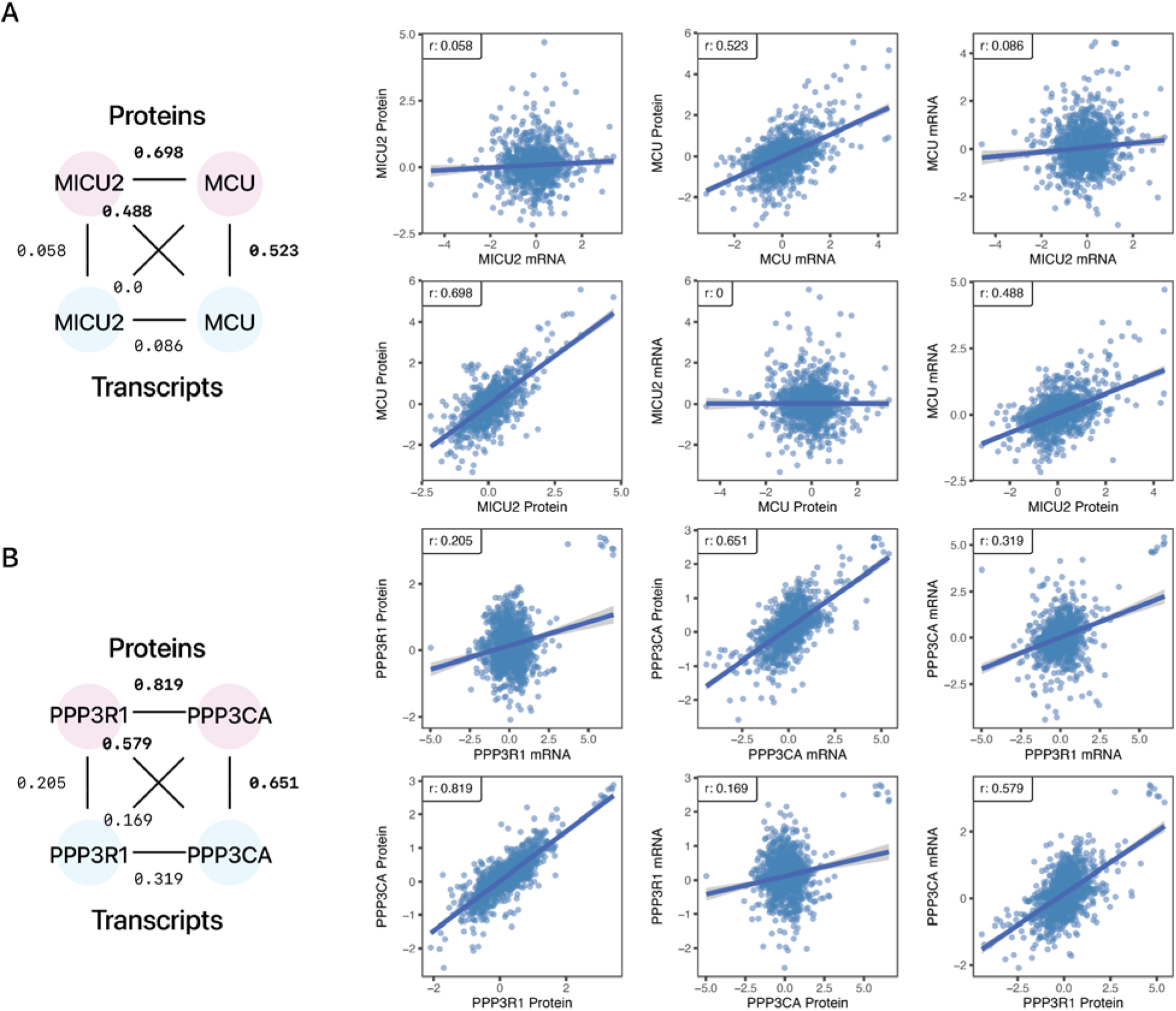
Cross-omics co-expression of MICU2 and PPP3R1 with functionally associated proteins. Two examples of proteins whose abundance is better explained by another transcript are shown. **A.** MICU2 protein level is predicted by MCU transcript but not its own transcript. **B.** PPP3R1 protein level is explained by PPP3CA transcript level but not its own transcript. Substantial correlations across transcripts and proteins (≥ 0.4) are bolded.

**Supplementary Figure S6:**
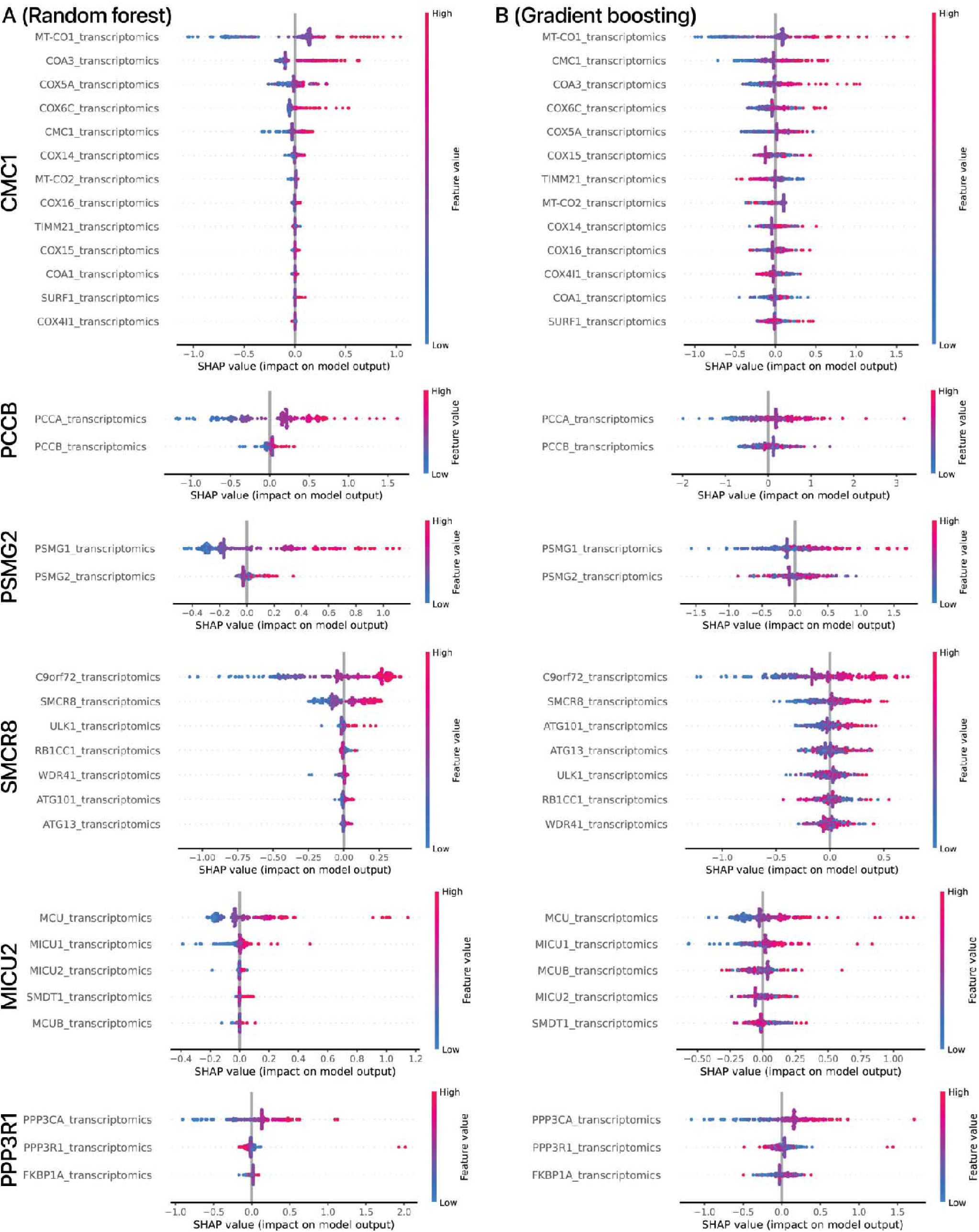
SHAP interpretation of feature importance in **A.** random forest and **B.** gradient boosting model output from the CPTAC_8 CORUM feature set. The SHAP values of top transcript features and their impact on model output are shown for each of six proteins highlighted in the text (from top to bottom): PCCB, CMC1, PSMG2, SMCR8, MICU2, PPP3R1.

**Supplementary Figure S7:**
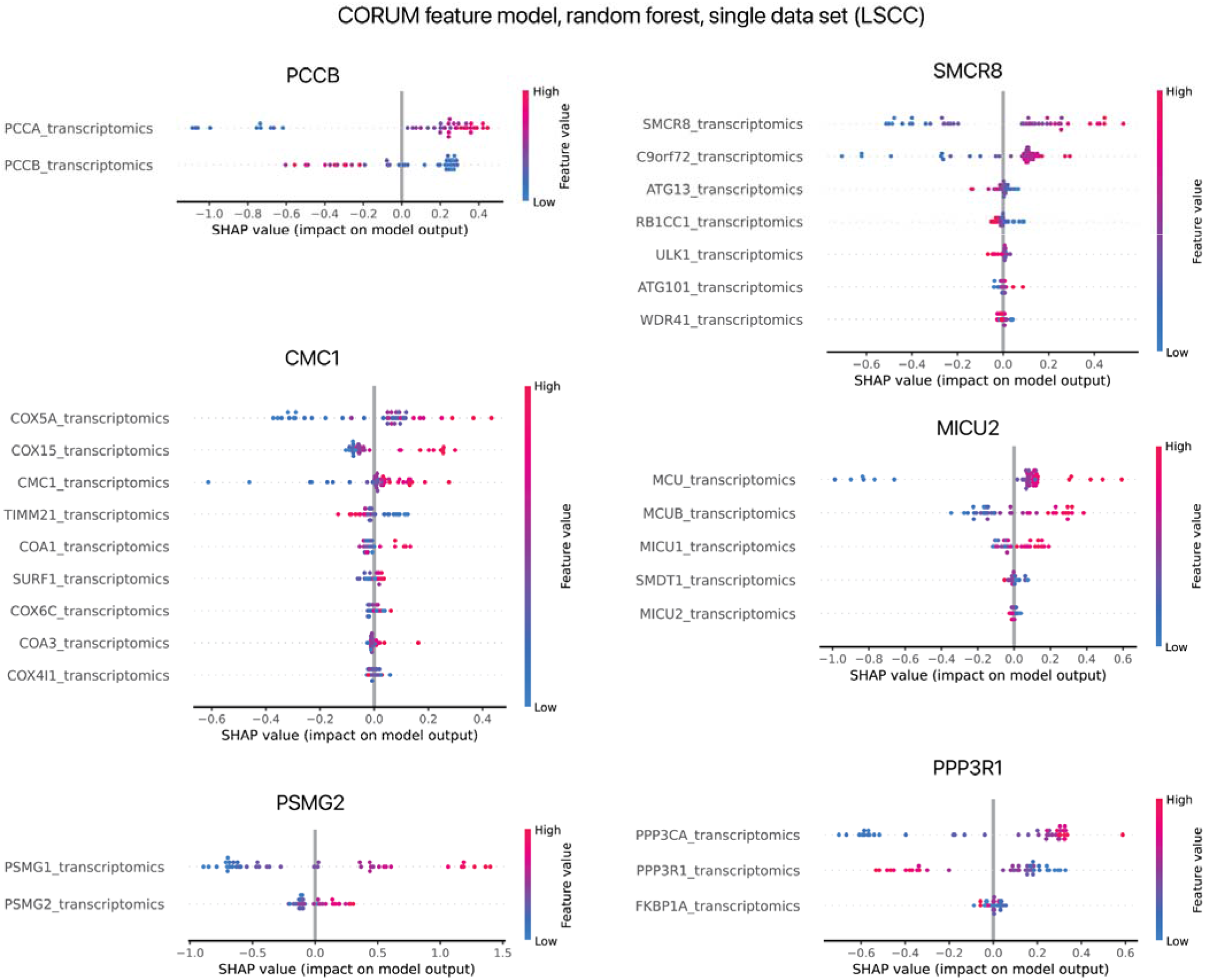
SHAP interpretation of feature importance in the CORUM feature set, random forest model, of a single data set (LSCC), showing largely conserved observations among the highlighted proteins PCCB, CMC1, PSMG2, SMCR8, MICU2, PPP3R1. Consistent with the analysis of the combined CPTAC_8 data set, five of the six proteins are not best predicted by their cognate transcripts and all but one (MT-CO1) of the top trans locus transcripts are preserved, which was not among the examined candidate features. Only test set data points are shown.

**Supplementary Figure S8:**
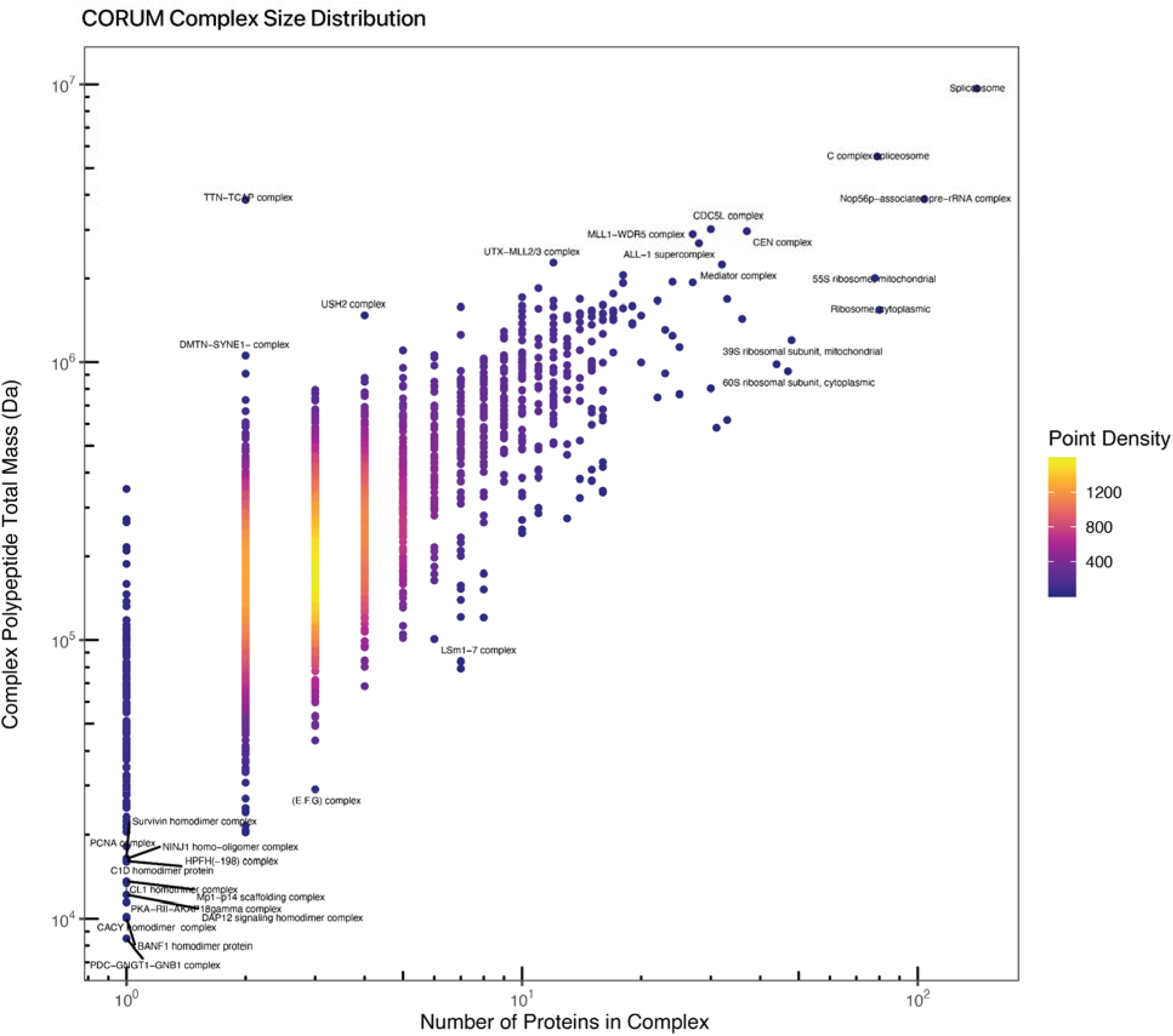
Density plot showing the distribution of complex sizes (x:-axis: number of member sin complex; y-axis: log10 total polypeptide molecular weight in complex) in the annotated feature set derived from CORUM v.3.0. Names of select complexes are labeled. The majority of complexes are small with a median of 3 proteins per complex.

**Supplementary Figure S9:**
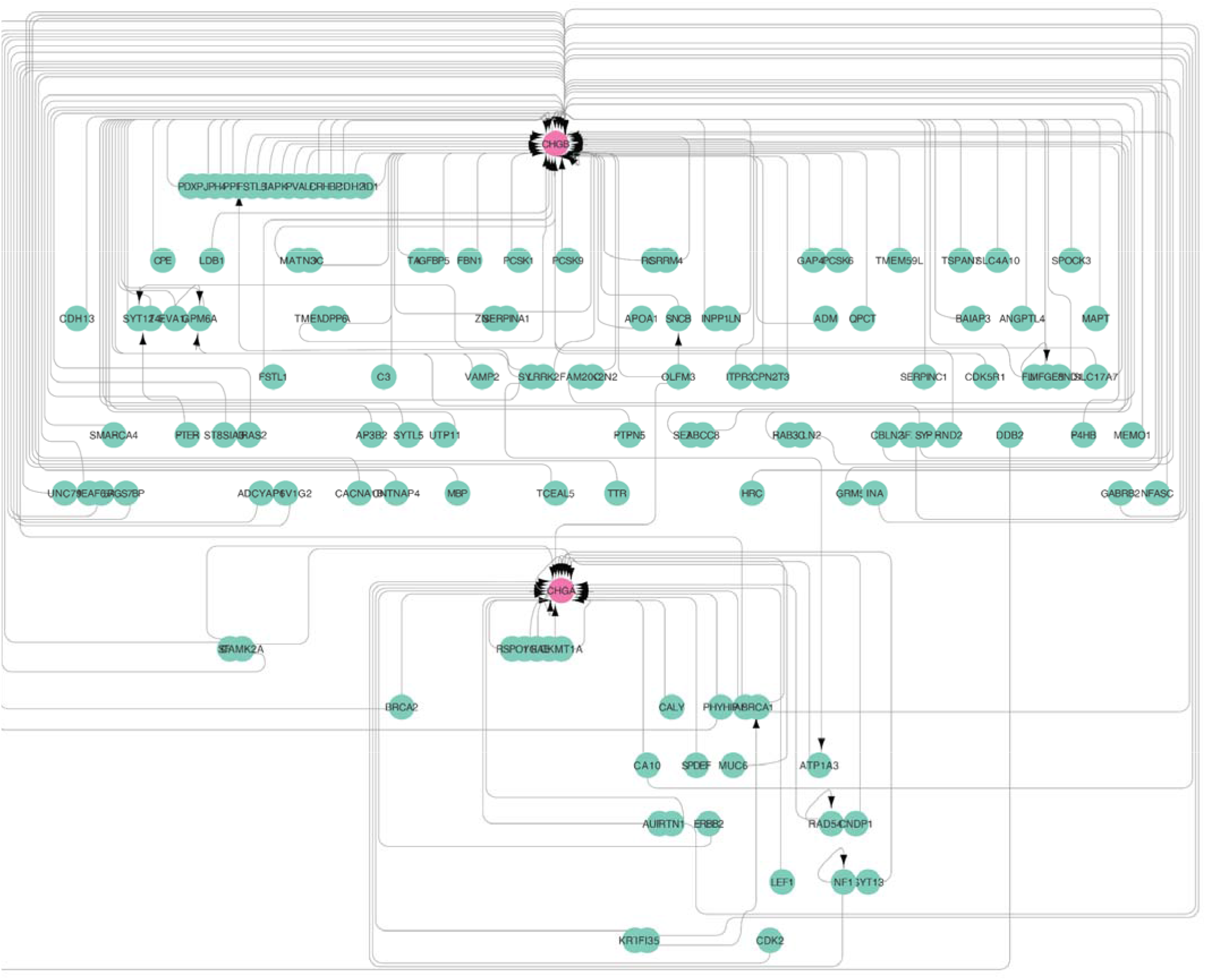
The most connected subgraph in the STRING feature set contains 6,319 nodes and 11,161 edges, containing proteins belonging to multiple distinct cellular compartments and multi-protein complexes. The hierarchical network diagram shows Chromogranin A and B (CHGA/CHGB) (magenta) and their first-degree inward flow neighbors (green) in the largest subgraph constructed from the STRING data set. The protein level of CHGA and CHGB is associated with the transcript levels of multiple secreted peptide coding genes, consistent with these proteins serve to regulate gene expression in secretion pathways.

## Supplementary Tables

Supplementary tables can be accessed on figshare at: https://doi.org/10.6084/m9.figshare.19330541

**Supplementary Table S1:** Table listing performance metrics (test set correlation coefficients, R^2^, NRMSE) of the tested models, feature sets, and data sets.

**Supplementary Table S2:** Table listing Gene Ontology cellular compartment term enrichment results for proteins that are well predicted by their transcript levels.

**Supplementary Table S3:** Table listing Gene Ontology cellular compartment term enrichment for proteins that are poorly predicted by their transcript levels.

**Supplementary Table S4:** Table listing individual genewise model performance metrics (test set correlation coefficients, R^2^, NRMSE) in the CPTAC_8 data set using elastic net models on different feature sets.

**Supplementary Table S5:** Table listing Gene Ontology cellular compartment term enrichment for proteins with substantial improvements in prediction performance upon incorporating STRING features in elastic net models.

**Supplementary Table S6:** Network edge list for proteins with improved predictions in the CORUM feature set along with the coefficients and feature importances of contributing transcripts.

**Supplementary Table S7:** Network edge list for proteins with improved predictions in the STRING feature set along with the coefficients and feature importances of contributing transcripts.

**Supplementary Table S8:** Hub genes based on centrality scores in the STRING feature set network.

